# Global and single-cell proteomics view of the co-evolution between neural progenitors and breast cancer cells in a co-culture model

**DOI:** 10.1101/2023.05.03.539050

**Authors:** Ole Vidhammer Bjørnstad, Manuel Carrasco, Kenneth Finne, Ingeborg Winge, Cecilie Askeland, Jarle B. Arnes, Gøril Knutsvik, Dimitrios Kleftogiannis, Joao A. Paulo, Lars A. Akslen, Heidrun Vethe

## Abstract

Tumor neurogenesis, a process by which new nerves invade tumors, is a growing area of interest in cancer research. Nerve presence has been linked to aggressive features of various solid tumors, including breast and prostate cancer. A recent study suggested that the tumor microenvironment may influence cancer progression through recruitment of neural progenitor cells from the central nervous system. However, the presence of neural progenitors in human breast tumors has not been reported. Here, we investigate the presence of Doublecortin (DCX) and Neurofilament-Light (NFL) co-expressing (DCX+/NFL+) cells in patient breast cancer tissue using Imaging Mass Cytometry. To map the interaction between breast cancer cells and neural progenitor cells further, we created an *in vitro* model mimicking breast cancer innervation, and characterized using mass spectrometry-based proteomics on the two cell types as they co-evolved in co-culture. Our results indicate stromal presence of DCX+/NFL+ cells in breast tumor tissue from a cohort of 107 patient cases, and that neural interaction contribute to drive a more aggressive breast cancer phenotype in our co-culture models. Our results support that neural involvement plays an active role in breast cancer and warrants further studies on the interaction between nervous system and breast cancer progression.

## Introduction

Tumor neurogenesis, the process by which nerves are formed and invade tumors, is a growing topic in current cancer investigations^1–3^. Increased nerve density is associated with tumor aggressiveness in several solid tumors, including breast, prostate, gastric, and head and neck cancers^4–8^. In breast cancer, presence of nerve fibers in the tumor microenvironment (TME) correlates with poor differentiation, a triple-negative phenotype, lymph node metastasis, and higher clinical stage^9–12^. Moreover, recent observations in prostate cancer mouse models have shown that the central nervous system (CNS) might influence cancer progression through recruitment of neural progenitor cells (NPCs) expressing doublecortin (DCX), revealing a novel mechanism for how nerves could be established in tumors^5^.

While NPCs might migrate to distant areas of the body, they arise from neurogenic zones in the adult CNS, *i.e.*, the subventricular zone near the lateral ventricles and the dentate gyrus of the hippocampus, where adult-born principal neurons are generated from adult neural stem cells (NSC)^13, 14^. NSCs can give rise to either NPCs or glial cells. Multipotent NPCs will differentiate into glial cells or immature CNS neurons that will integrate into pre-existing neural networks in the brain^15^.

It is not known whether human breast tumors contain NPC-like cells, or how NPCs interact with breast cancer cells. Obtaining CNS-derived neurons from living humans that maintain a differentiation potential for *in vitro* studies is challenging. However, stem cell technology and differentiation of pluripotent stem cells into specific human cell populations represents an impactful tool for developmental studies, disease modeling and regenerative medicine^16, 17^. Such technology can be used to generate otherwise quite inaccessible human cell populations for *in vitro* models.

Here, we aim to explore the presence and spatial localization of DCX+/NFL+ cells in primary breast cancer tissue by Imaging Mass Cytometry (IMC). We generate an *in vitro* model of breast cancer innervation, and capture the mutual interactions and co-evolution of breast cancer cells and neural cells in co-culture by mass spectrometry proteomics.

## Results

### Breast tumors show presence of cells expressing the neural progenitor marker doublecortin

DCX is a marker associated with neural progenitor cells and the axonal growth cone of migrating central and peripheral neurons^18^, and has been used to detect neural recruitment processes. Recent findings in mouse models of prostate and breast tumors have demonstrated that DCX+ neural progenitors migrate from the central nervous system (CNS) to primary and metastatic tumor sites. After migration, these DCX+ cells in the TME will differentiate to form maturing neurons contributing to tumor progression^5^. To test whether human primary breast tumors contain DCX-expressing cells, we used immunofluorescence (IF) staining for DCX in basal-like and luminal-like breast tumors (Fig. 1a, b). We found positive stromal staining for DCX, similar to the findings in prostate tumors^5^. DCX+ cells did not express markers of mature peripheral nerve fibers (Neurofilament-Heavy (NFH)).

**Figure 1.**
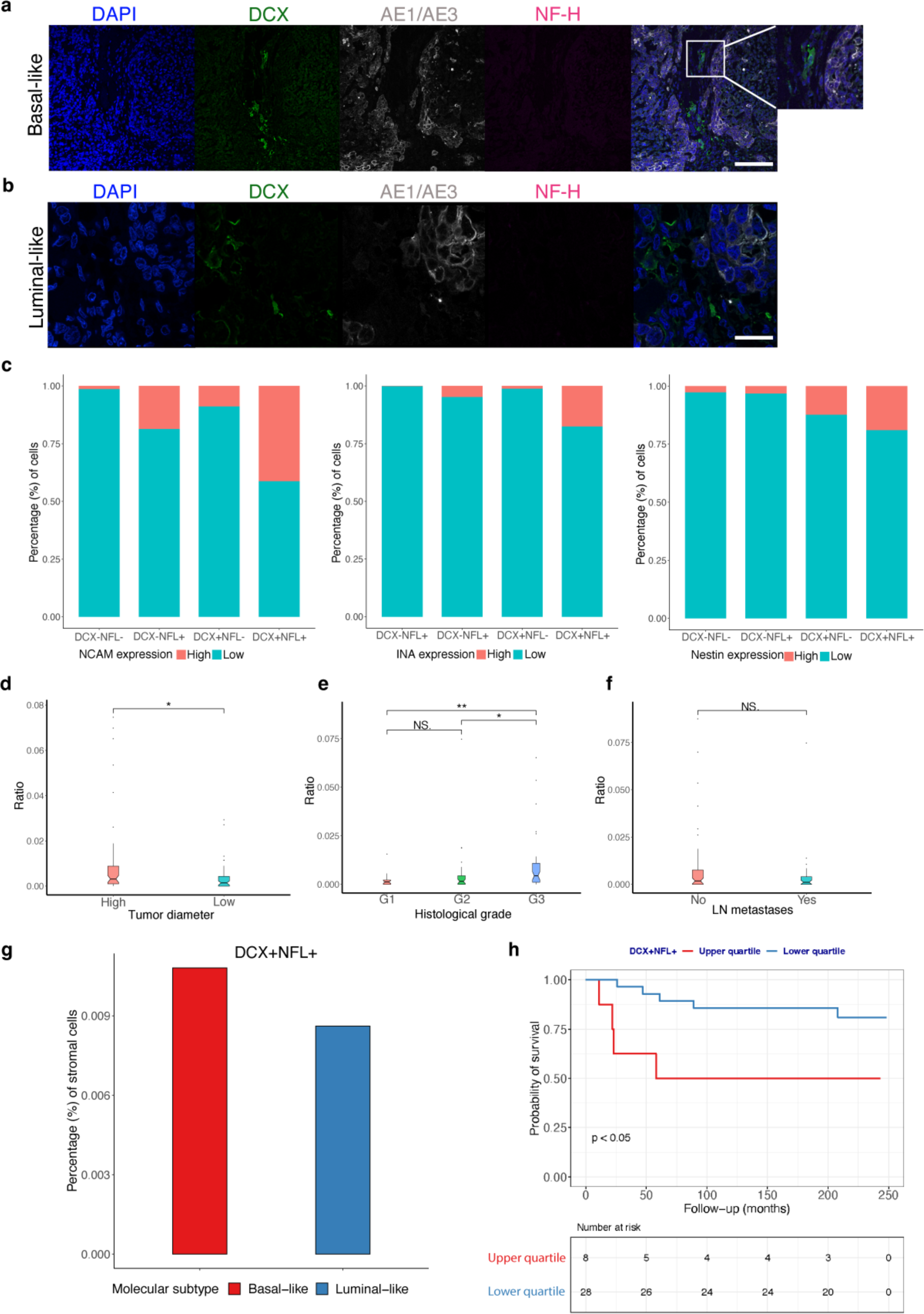
Doublecortin is expressed in the breast cancer microenvironment. **a** Confocal microscope pictures of basal-like and **b** luminal-like breast cancer tissue. DCX expression (green), AE1/AE3 (grey), Neurofilament-Heavy (NFH, negative). **c** Population distribution of DCX^+^/NFL^+^ cell populations for the markers NCAM, INA and Nestin. **d** Tumor diameter difference (significance was determined by Mann–Whitney U test (*p = 0.028)) by presence of DCX^+^/NFL^+^ cell populations in breast cancer. **e** Histological grade (significance was determined by Mann–Whitney U test (*p = 0.015 **p = 0.003)) by presence of DCX^+^/NFL^+^ cell populations in breast cancer. **f** Lymph node metastasis by presence of DCX^+^/NFL^+^ cell populations in breast cancer (p=ns). **g** Proportion of stromal DCX+/NFL+ in basal-like (number of cells: n=671 DCX+/NFL+ cells/ n=62071 all cells analyzed) and luminal-like (number of cells: n=465 DCX+/NFL+ cells/ n=5398 all cells) breast cancer tissue. **h** Kaplan-Meier patient survival plot by content of DCX^+^/NFL^+^ cell populations. Number of patients is displayed under the plot. (Significance was determined by the log-rank test (*p = 0.023)). Scale bar A – 100 μm B – 42 μm.

To investigate the population of stromal DCX+ cells present in breast tumors in more detail using additional markers by IMC, we designed a panel of 38 antibodies, containing DCX, NFL, Nestin, Neural Cell Adhesion Molecule (NCAM) and Internexin-alpha (INA) as representative neural markers (Supplementary Table 1)^19^. To confirm a neural phenotype of the cells expressing DCX, we focused on cells that were double positive for DCX and NFL, identifying a small population of 7,930 DCX+/NFL+ cells among 413,271 single-cells or equally denoted cellular units (1,96 % of the total cell population) in tissue from 107 breast cancer cases, including 56 luminal-like (luminal A n=26, luminal B n=30) and 52 with triple-negative (basal- like) subtype (Supplementary Table 2). Further investigations of this population show that they also express other markers of NPCs including Nestin, NCAM and INA, as shown by higher expression of these markers as compared to all other cells (Fig. 1c), supporting a neural phenotype of the DCX+/NFL+ cells.

To assess the spatial localization of DCX+/NFL+ cells, we first categorized our IMC single-cell data into a tumor cell compartment (marked by AE1/AE3) and a stromal compartment (marked by vimentin and aSMA) within the luminal-like and basal-like breast cancer subtypes. By analyzing the full cohort of 107 cases, we found that the stromal compartment contained 1% DCX+/NFL+ cells.

### Double expression of DCX and NFL correlates with tumor aggressiveness in breast cancer

To assess the potential clinical relevance of neural progenitors in breast cancer, cells co-expressing DCX and NFL were quantified across our cohort of 107 breast tumors (Supplementary Figure 1). The presence of DCX+/NFL+ cells in the stromal compartment was significantly associated with tumor aggressiveness, as shown by higher number of DCX+/NFL+ cells in breast tumors with large tumor diameter (Fig. 1d) and higher histological grade (Fig. 1e). No significant difference with regards to lymph node metastasis was found (Fig. 1f). When comparing the proportion of stromal DCX+/NFL+ cells in basal-like versus luminal-like breast cancer tissue, we found higher numbers of DCX+/NFL+ cells in basal-like breast tumors (Fig. 1g, Supplementary Table 3). Based on our size limited cohort, higher number of DCX+/NFL+ cells in breast tumor stroma were associated with shorter patient survival (Fig. 1h). In contrast, the presence of DCX^+^/NFL^+^ cells in the epithelial compartment showed no significant correlation with tumor diameter, histological grade, lymph node metastasis, or patient survival. Our data indicates that stromal presence of DCX+/NFL+ cells may play a role in breast cancer progression which warrants further investigations into the direct relationship between neural progenitors and breast cancer cells.

### Human pluripotent stem cells give rise to neural progenitor cells

To investigate the direct interaction between relevant NPCs from the CNS and breast cancer cells, we generated NPCs from the human induced pluripotent stem cell (hiPSC) line XCL-1 DCXp-GFP and human embryonic stem cells (hESCs) H9inGFPhESCs. To test the differentiation potency of NPCs into mature neuron-like cells *in vitro*, we differentiated the XCL-1 DCXp-GFP line based on a well-established differentiation protocol generating dopaminergic neurons^20^. Our stem cell-derived NPCs are proliferative (marked by Ki67) and express neural markers such as: DCX, INA, NeuN, Nestin and NFH (Fig. 2a). Following *in vitro* differentiation for up to three weeks, our maturing neurons express tyrosine hydroxylase (TH). We subsequently differentiated H9inGFPhESCs towards NPCs, midbrain dopaminergic neurons and matured these for two and five weeks.

**Figure 2.**
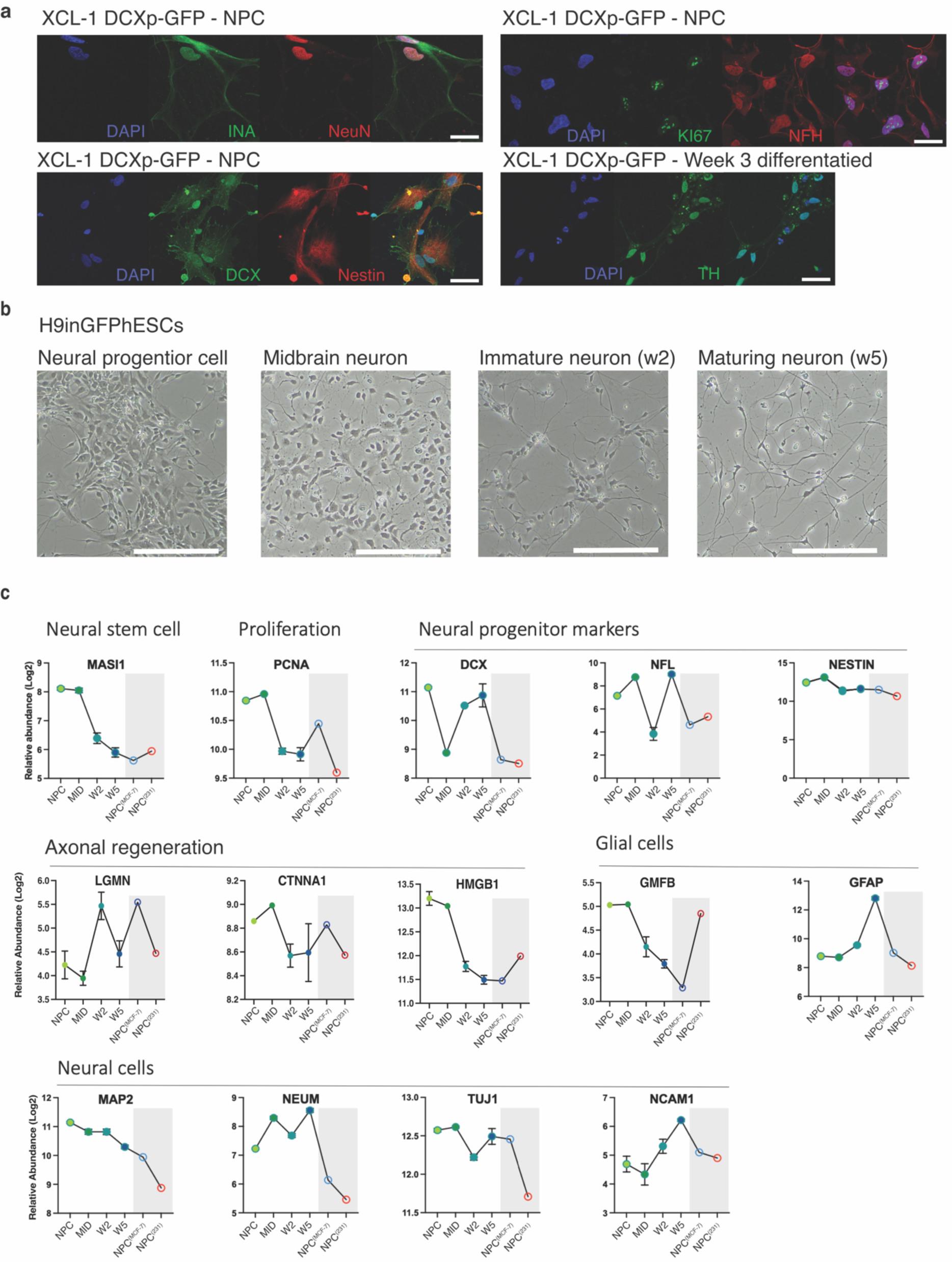
Neural progenitor cell differentiation. **a** Sp8 confocal images showing the marker presence of INA, NeuN, DCX, Nestin, Ki67, NFH and TH in the XCL-1 DCXp-GFP cell line. Scale bar – 40 μm **b** Brightfield images displaying the morphological change in the H9inGFPhESCs upon differentiation into NPC and subsequent differentiation into midbrain dopaminergic neurons, immature neurons (after two weeks of differentiation) and maturing neurons (after five weeks of differentiation). Scale bar – 200 μm. **c** Selected protein abundances of neural markers following the differentiation steps of NPCs, midbrain neurons, immature neurons, maturing neurons, NPC after co-culture with MDA-MB-231 spheroids and with MCF7 spheroids.

Based on the ability of XCL-1 DCXp-GFP NPC to differentiate into TH^+^ neurons (Fig. 2a). We aimed to investigate changes in the differentiation potential of NPCs upon interaction with breast cancer cells, we tested *in vitro* NPC differentiation of H9inGFPhESCs cells towards midbrain-patterning neural precursors and maturing dopaminergic neuron-like cells (Fig. 2b). Characterization of cells was based on protein profiles specific to neural stem cells, neural progenitor cells, astrocytes and maturing neural cells. As expected, the protein profile of the neural stem cell marker (Mushi 1) and proliferation marker (PCNA) decreased during differentiation and subsequent maturation (Fig. 2c). Our hESC-derived NPCs also expressed DCX, the neural growth cone marker neuromodulin (NEUM)^21^, other neural markers including Nestin, NFL, NCAM and TUJ1 and markers of axonal regeneration (LGMN, CTNNA1, HMGB). We found an increased expression of the astrocyte marker glial fibrillar acidic protein (GFAP) following *in vitro* maturation, whereas the glial maturation factor beta (GMFB) decreased. This suggests a heterogeneous cell population at the end of differentiation containing both neural and glial cells, confirming the differentiation potential of NPCs.

### Neural progenitor cells interact with breast cancer cells in 3D models *in vitro*

To model breast cancer innervation, we generated breast cancer spheroids from three basal-like (MDA-MB-231, HS578T, BT549) and three luminal-like (MCF-7, BT474, T47D) breast cancer cell lines, and combined these with H9inGFPhESC-derived NPCs in co-culture (Fig. 3a). Our co-cultures were maintained for seven days; at day six, doxycycline was added to the media for induction of GFP expression in NPCs (Fig. 3b). At day seven, co-cultured spheroids were collected for whole mounting or cell sorting (Fig. 3c) and subsequent downstream proteomics analysis. Our basal-like breast cancer spheroids (MDA-MB-231 and BT549) showed a typical grape-like morphology, with less densely packed spheroids as compared to HS578T, and our luminal-like breast cancer spheroids (MCF-7, BT474, T47D), which showed a mass-like morphology^22^, displaying a more distinct cell-cell adhesion forming more aggregated morphology (Fig. 3d-e). The spheroids varied in diameter, co-cultures of MDA-MB-231+NPC, BT549+NPC and HS578T+NPC grew to around 3-500 μm. HS578T+NPC co-cultures were more spherical, assuming a circumscribed and firm morphology. MCF-7+NPC and BT474+NPC formed tight spheroids containing cystic pockets, with an average diameter of 100-250 μm, where BT474+NPC formed larger spheroids. Smaller T47D+NPC spheroids seem to fuse together and formed densely packed spheroids up to 1500 μm in diameter, with no visible cystic areas. When comparing breast cancer spheroids before and after co-culture with NPCs, no discernible morphological differences were observed.

**Figure 3.**
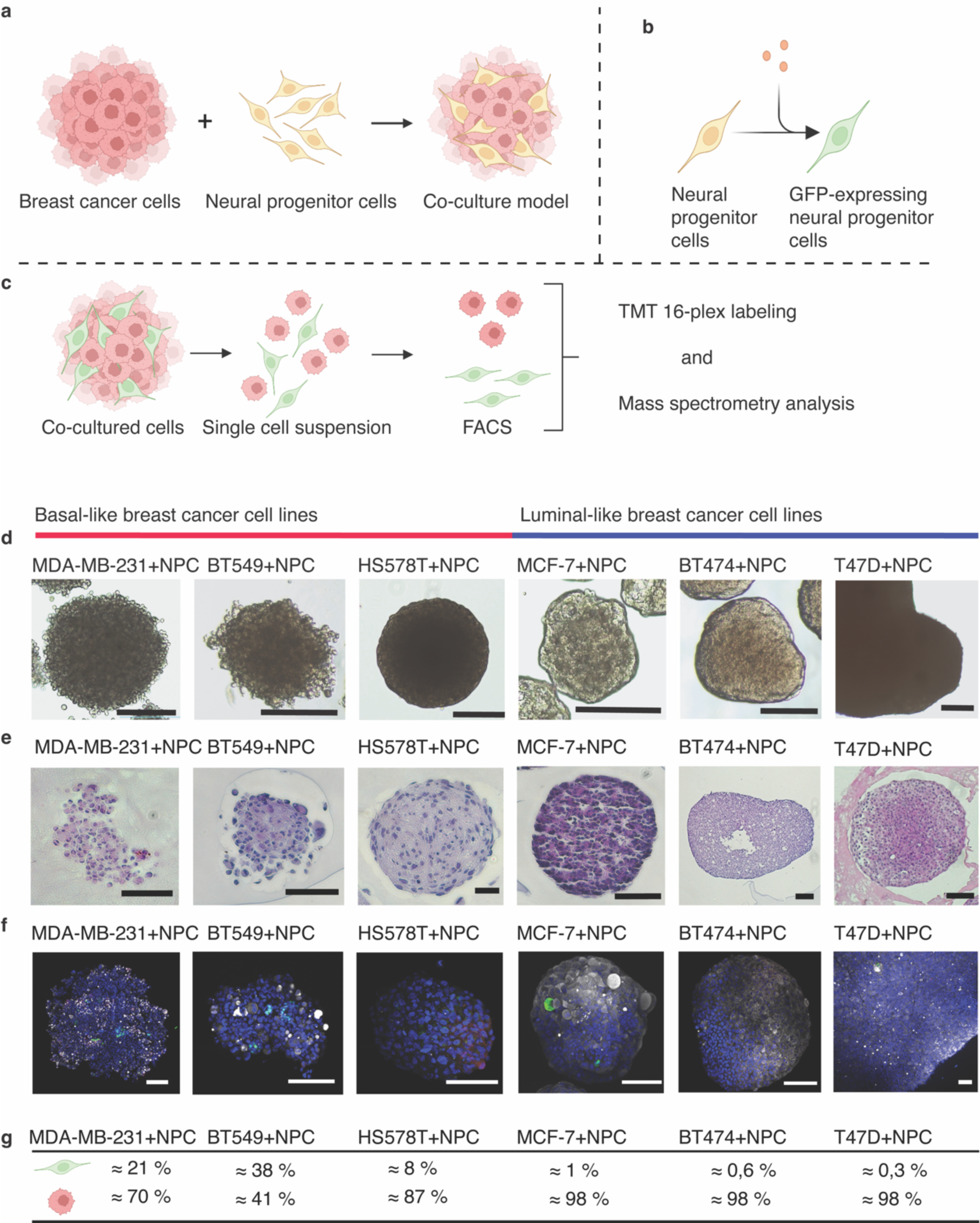
Model setup of co-cultured breast cancer cell lines and NPCs. **a** Schematic presentation of our co-culture model, where breast cancer cells were combined with NPCs to form a 3D co-culture model. **b** GFP-expression induction by doxycycline in hESCs-derived NPCs. **c** Spheroids containing GFP-expressing NPCs and cancer cells are disaggregated into a single-cell suspension. This single cell suspension was then FACS sorted. Cells both negative and positive for GFP are subsequently prepared and conducted for mass spectrometry analysis. **d** Brightfield photos of spheroids when combining breast cancer cell lines and NPCs. **e** H&E staining of the respective cell lines. **f** High magnification immunofluorescence photos following whole mounted immunofluorescent on cancer-NPC spheroids (DAPI – blue, GFP – green, DCX – red, AE1/AE3 - grey). **g** FACS distribution of GFP positive events and negative for spheroid sorting. Scale bars: d = 250 μm, e and f = 100 μm.

NPCs integrated well with the basal-like MDA-MB-231 and BT549 spheroids, where co-cultures contained multiple groups of NPC cells (Fig. 3f), as shown by FACS sorting: 20 % and 37% GFP+ cells, respectively (Fig. 3g & Supplementary figure 2). Fewer NPCs (8%) were integrated in the HS578T+NPC spheroids. For the luminal-like spheroids, MCF-7 integrated NPC cells best, however, compared to basal-like this integration was marginal at around 1-2%, while FACS sorting of GFP-expressing NPCs from BT474 and T47D spheroids ranged from 0,3-0,6%. Most luminal-like spheroids seem to integrate a couple of individual cells, but not at similar levels as basal-like breast cancer spheroids, suggesting that basal-like breast cancer cells have a stronger attraction for NPCs than luminal-like breast cancer cells.

### Global proteome changes in breast cancer cells after interaction with NPCs

The interaction between tumor cells and nerves is bidirectional, as cancer cells can secrete neurotrophic factors, neurotransmitters, and axon guidance factors to stimulate nerves sprouting and infiltration, while nerves can secrete neuroactive factors that can boost tumor growth and spread^4, 23, 24^. To characterize the effect of NPC interaction on the breast cancer cellular proteome, we compared the proteome of FACS-sorted breast cancer cells from the three basal-like cells (MDA-MB-231, HS578T, BT549) and three luminal-like (MCF-7, BTB474, T47D) spheroid models before and after co-culture with GFP-expressing NPCs. Overall, our global proteomics comparison showed that NPC-interaction changed the expression pattern of 3643 proteins in breast cancer cells (BH adjusted FDR < 0.05). Hierarchical clustering, multidimensional scaling (MDS) and correlation analysis indicated a clear separation of breast cancer cells before and after NPC interaction (Fig. 4a-c), except for HS578T^(NPC)^ which positioned near HS578T, which could indicate different degrees of NPC-integration with breast cancer cells (as shown in Fig.3).

**Figure 4.**
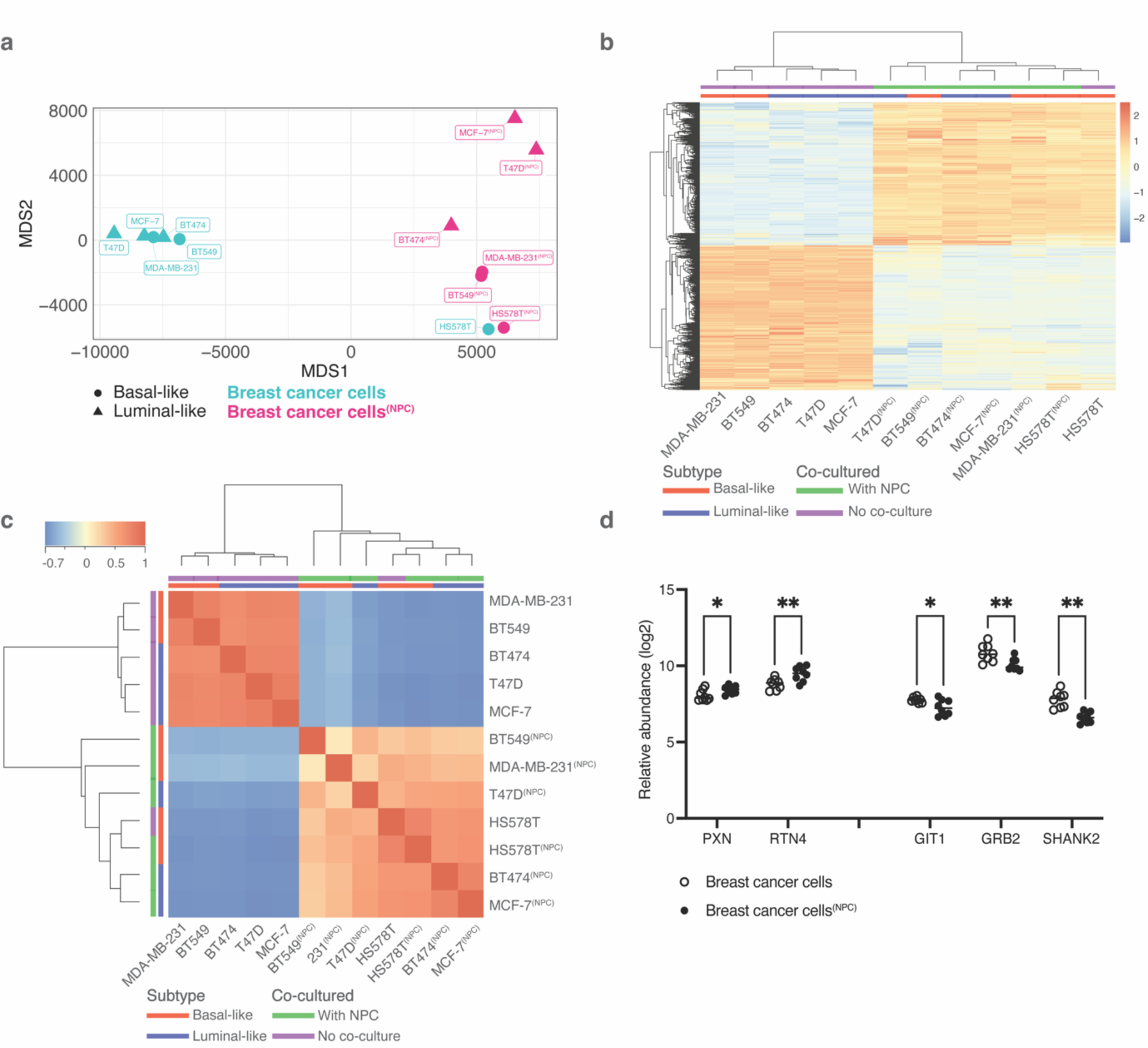
Proteomic landscape before and after co-culture. **a** Multidimensional scaling plot displaying the sample proteome after the selection of statistically significant proteins before and after co-culture with NPCs (Benjamini Hochberg (BH) adjusted FDR < 0.05). Superscripted text indicates the co-cultured condition. **b** Hierarchical clustering of normalized TMT-ratios pooling all breast cancer cell lines together and comparing statistically significant proteins before and after co-culture with NPCs (BH adjusted FDR < 0.05). Color bar: Blue represents a downregulated protein expression compared to the mean intensity with a gradient into beige for no change in expression and into red for upregulated expression. **c** Correlation plot of expression between cell lines pooled all breast cancer cell lines together and comparing statistically significant proteins before and after co-culture with NPCs (BH adjusted FDR < 0.05). Each square represents the Spearman correlation coefficient between two samples with comparisons being hierarchically clustered as well. Color bar: Blue represents a negative correlation with a gradient into beige for no correlation and into red for positive correlation. **d** Selected proteins involved in axon guidance with changed expression pattern before and after co-culture with NPCs (*P ≤ 0.05, **P ≤ 0.01, significance was determined by unpaired t test).

### Co-culture with NPCs induces aggressive tumor features in breast cancer cell lines

With a statistical overrepresentation test using PantherDB, we found that 10% of the proteins with altered expression in breast cancer cells were involved in axon guidance, nervous system development, and signaling via ROBO receptors (232 proteins, FDR 3.92E-23, Fisher’s Exact Test (Supplementary Table 4)). To further investigate whether NPCs induced aggressive tumor features in breast cancer spheroids, we explored the expression profile of proteins involved in *axon guidance*, *epithelial-to-mesenchymal transition (EMT)*, *migration*, *invasion*, and *proliferation*. Axon guidance signaling proteins Paxillin (PXN) and Reticulon4 (RTN4) showed increased abundance, while GIT1, GRB2 and SHRANK2 showed decreased abundance in breast cancer cells after co-culture (Fig. 4d) (Supplementary Table 5).

EMT is considered a major mechanism responsible for breast cancer progression^25^. Since EMT-associated features are expressed differently in basal-like and luminal-like cell lines^26^, we performed an additional TMT 16-plex experiment focusing on one basal-like cell line (MDA-MB-231, n=3) and one luminal-like cell line (MCF-7, n=3) before and after co-culture with NPC (Fig. 5). When comparing breast cancer spheroids before and after NPC interaction, we found 946 differentially expressed proteins (DEPs) in the basal-like breast cancer MDA-MB-231 spheroids (FC >±1.5, Benjamini Hochberg (BH) adjusted FDR < 0.01) (Fig 5a, c), while luminal-like MCF-7 spheroids showed 787 DEPs (FC >±1.5, BH adjusted FDR < 0.01) (Fig. 5b, c). Analyzing the upstream regulators after co-culture, both MDA-MB-231^(NPC)^ and MCF-7s^(NPC)^ top three regulated proteins were TP53, MYC and HNF4A (Fig. 5d, Fig. 5e), with MAPT and ESR1 being the subsequent ranked upstream regulators for MDA-MB-231^(NPC)^ and CLPP and GABA for MCF-7^(NPC)^. Network analysis of the comparative analysis displayed the key predicted master regulators: RIPK2, CEBPD and FASN (Fig. 5f). We found the expression profile of key regulators predicted activation of processes such as “*cell proliferation of tumor cell lines*”, “*invasion of tissue*”, “*metastasis*” and “*migration of tumor cells lines*” for both MDA-MB- 231^(NPC)^ and MCF-7^(NPC)^, suggesting that NPC interactions with our two breast cancer cell lines induced aggressive tumor features. Comparative analysis between MDA-MB-231(NPC) and MCF-7(NPC) breast cancer cell lines following co-culture predicted that the upstream regulators most affected by NPC interaction showed similar alteration levels in both cell lines (Fig. 5g). This included stem cell markers such as MYC, POU5F1 (OCT4), LET-7 (Lin28), the microenvironmental stress response marker NUPR1, and the neuronal differentiation marker HMG20A, which together may serve as a panel of candidate markers for further evaluation of neural interaction in other breast cancer cell lines.

**Figure 5.**
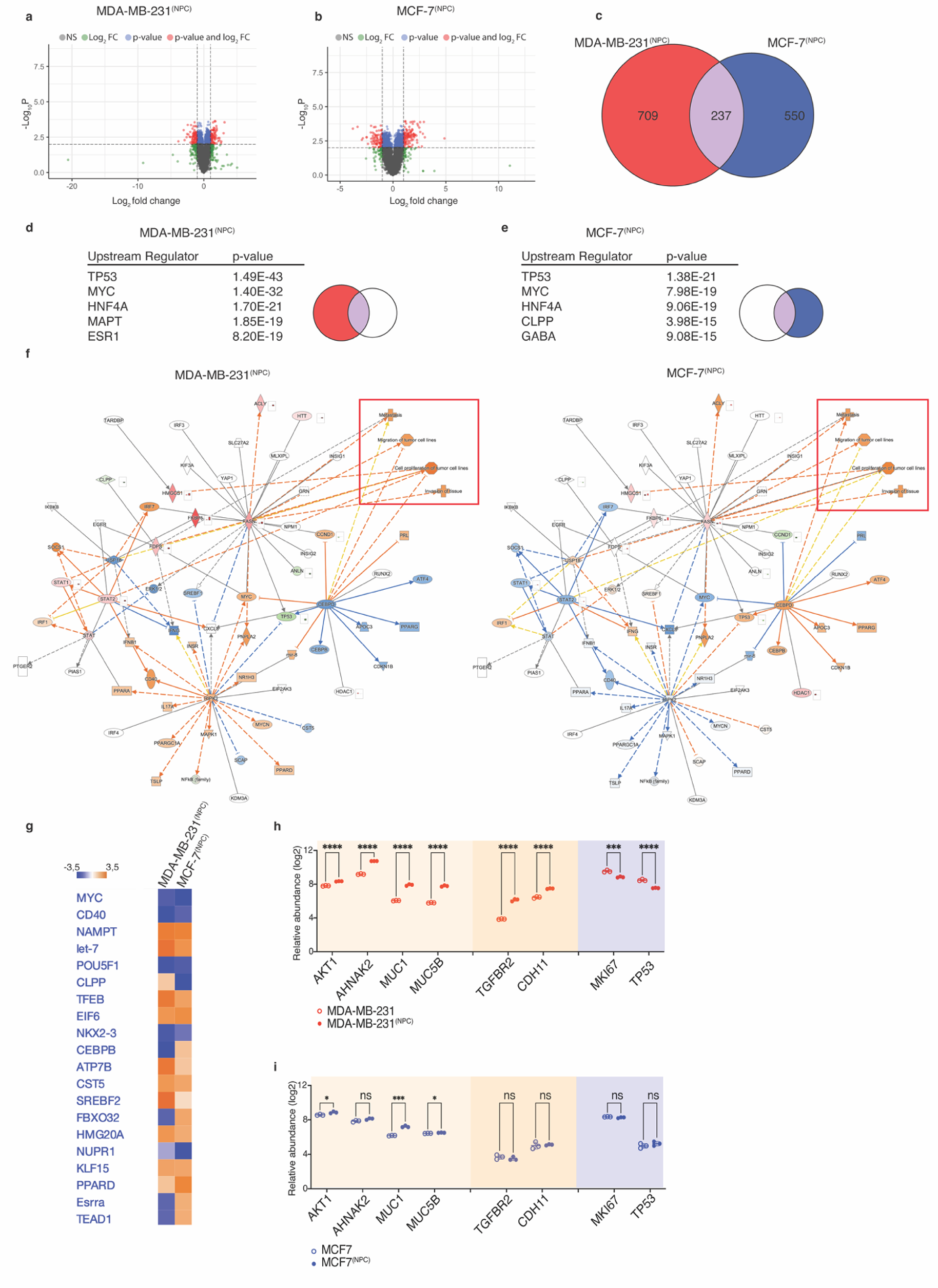
Differentially expressed proteins (DEPs) for MDA-MB-231 and MCF-7 after co-culturing with NPCs. **a** Volcano plot representing DEPs for MDA-MB-231^(NPC)^ relative to MDA-MB-231 (BH adjusted FDR < 0.01, fold change in gene expression > 1.5), from which 946 DEPs were quantified. **b** Volcano plot representing DEPs for MCF-7^(NPC)^ relative to MCF-7 (BH adjusted FDR < 0.01, fold change in gene expression > 1.5), from which 787 DEPs were quantified. **c** Venn diagram representing the common DEPs between MDA-MB-231^(NPC)^ and MCF-7^(NPC)^. MDA-MB-231^(NPC)^ had 709 unique DEPs and MCF-7^(NPC)^ had 550, with 237 common. **d** IPA generated graphical representations of predicted upstream regulators by DEPs for MDA-MB-231^(NPC)^ with associated p-values (Red and purple, left and middle of Venn diagram.) **e** IPA generated graphical representations of predicted upstream regulators by DEPs for MCF-7^(NPC)^ with associated p-values (Purple and blue, middle, and right of Venn diagram). **f** Selected radial network for the up- and downregulated proteins by the common DEPs between MDA-MB- 231^(NPC)^ and MCF-7^(NPC)^. Downstream effects are based on preexisting IPA protein lists. The red rectangle indicates the predicted activated processes of “Metastasis”, “Migration of tumor cell lines”, “Cell proliferation of tumor cell lines” and “Invasion of tissue” for both MDA-MB-231^(NPC)^ and MCF-7^(NPC)^. Orange – predicted activation, blue – predicted inhibition, red – increased measurement, green – decreased measurement. **g** IPA generated upstream analysis showing upstream regulators of comparison analysis for MDA-MB-231^(NPC)^ and MCF-7^(NPC)^. h Relative abundance levels based on quantitative proteomics measurements of single-protein levels of markers involved in EMT, cell migration and proliferation in MDA-MB-231^(NPC)^ cells and i MCF- 7^(NPC)^ cells (not significant (n.s.) P>0.05, *P ≤ 0.05, ***P ≤ 0.001, ****P ≤ 0.0001, significance was determined by unpaired t test).

To investigate further the predicted aggressive phenotypic alterations observed, we also investigated the expression of specific markers: EMT markers such as AKT1, AHNAK2, MUC1 and MUC5B were upregulated in both cancer cell lines (Supplementary Table 6 & 7). In MDA-MB-231^(NPC)^, we detected a significant upregulation of TGFBR2 and CHD11 (Fig. 5h), whereas the expression levels of these two proteins were not significantly changed in MCF-7^(NPC)^ (Fig. 5i), reflecting the impact of subtype differences between basal- and luminal-like cell lines. The proliferation marker Ki67 was significantly decreased in breast cancer spheroids after co-culture with NPCs. Finally, the top predicted protein TP53 was downregulated in MDA-MB- 231^(NPC)^ (Fig. 5h). These findings indicate that breast cancer cell lines are affected by the interaction with NPCs towards a more aggressive phenotype, as shown both by predicted activation of biological processes and by single-protein markers for proliferation, TP53 and EMT, with significant abundance differences in MDA-MB-231.

### Co-culture with breast cancer cells leads to activation of neurogenesis in neural progenitors

Neural progenitor cells isolated from prostate tumors in mouse models, have previously been shown to induce intratumor neurogenesis and differentiate into mature sympathetic neurons ex vivo5. To explore how co-culture with breast cancer cells affected the NPC proteome, we compared sorted NPCs before and after co-culture with breast cancer spheroids and found 235 differentially expressed proteins (Supplementary Table 8, FC >±1.5, BH adjusted FDR < 0.05). Network analysis was conducted to determine potential upstream regulators responsive for driving the proteomics changes in NPCs (Fig. 6a). Comparing effects of the upstream regulator network we found biological processes such as “*neurogenesis*” and “*development of nervous tissue*” predicted to be activated, while the term “*proliferation of neural cells*” was predicted to be inhibited. Considering the top three canonical pathways affected, remodeling of epithelial adherence junctions was the most significant change, with altered BAG2 and FAT10 signaling following, displaying an alteration in cellular adhesion and proteasomal degradation (Fig. 6b). The top upstream regulators were MAPT, TP53, APP, AYC and PTP4A1. Neuronal proteins such as DCX, NFL, STMN1 and GABARAP were all downregulated after co-culture with breast cancer cells (Fig. 6c).

**Figure 6.**
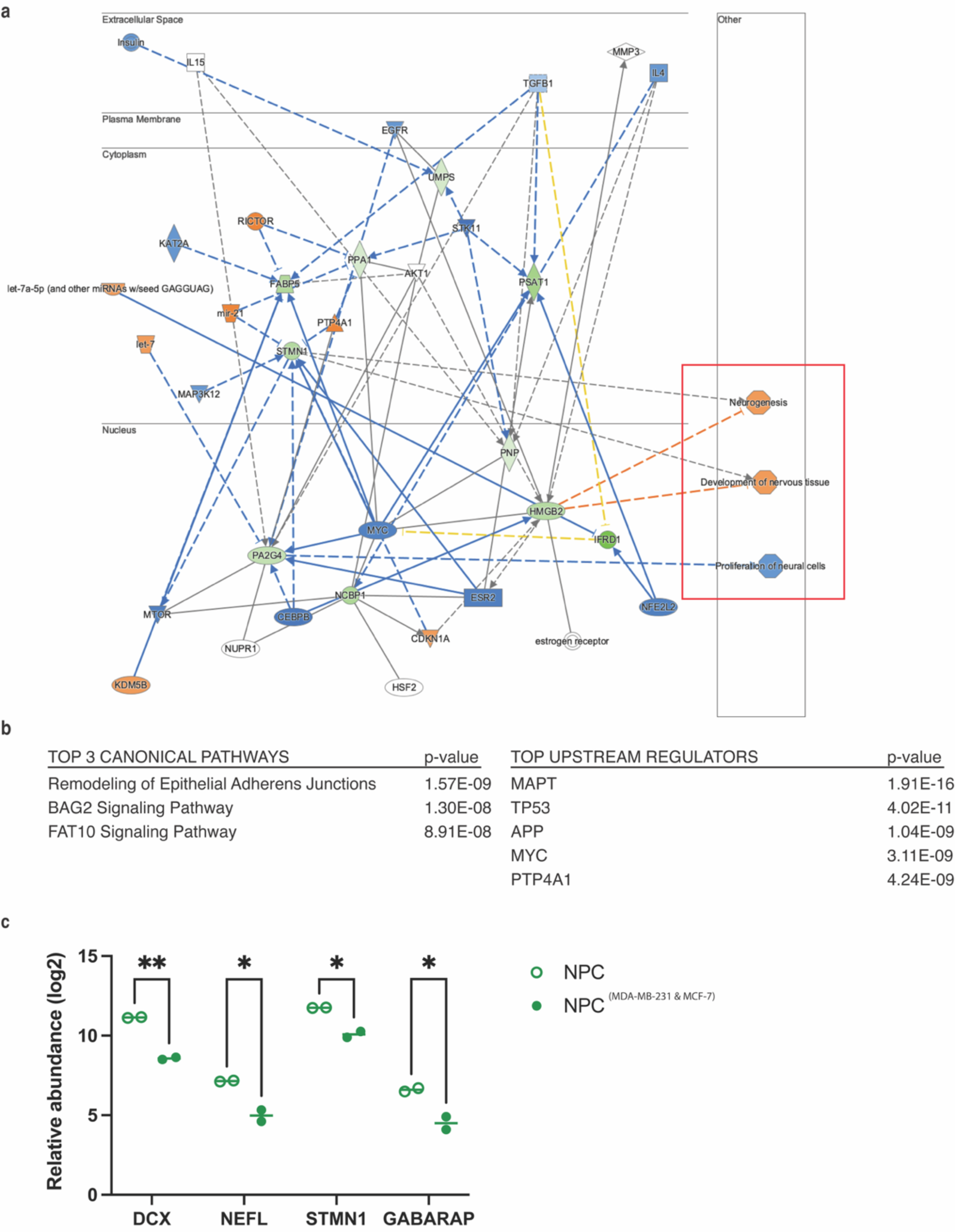
Pathway analysis of neural progenitor cells after co-culture with breast cancer cells. **a** IPA generated graphical representations of predicted upstream regulators and their downstream effects. NPC samples are compared to the average protein counts of NPC^(MDA-MB- 231 & MCF-7)^ (BH adjusted FDR P < 0.05). In total, 787 DEPs were quantified. Downstream effects are based on preexisting IPA protein lists. The red rectangle indicates the downstream activated processes of “Neurogenesis”, “Development of nervous tissue” and “Proliferation of neural cells”. Orange – predicted activation, blue – predicted inhibition, red – increased measurement, green – decreased measurement. **b** Top 3 canonical pathways and top upstream regulators effected by co-culturing with breast cancer cell lines of NPCs compared to the average protein counts of NPCs^(MDA-MB-231 & MCF-7)^. **c** Selected neural markers were lower expressed in the NPC^(MDA-MB-231 & MCF-7)^ compared to NPC in culture (*P ≤ 0.05, **P ≤ 0.01, significance was determined by unpaired t test).

Overall, our data indicate that stromal presence of DCX+/NFL+ cells in breast tumors (n=107) associates with aggressive tumor features, suggesting that these cells may play a role in breast cancer progression. Our *in vitro* model with NPC integration in breast cancer spheroids allowed us to monitor how these cells interact in real time and revealed changes in the proteome of breast cancer cells as they co-evolve during interaction with NPCs and identified key regulators of this communication.

### Discussion

Cancer neuroscience is an emerging research field with discoveries relating to the long-overlooked role of nerves in cancer^1, 2^. An increasing number of studies have shown nerve fiber involvement in cancer progression^4, 6, 27, 28^. Most reports have focused on the role of peripheral nervous system by axonogenesis (the outgrowth of axons from pre-existing nerves found in the TME^27^). However, recent observations in prostate cancer, indicate a new route of cancer-neural cell crosstalk coming from CNS-derived neural progenitor cells marked by DCX^5^. In our present study, we explored whether a similar process also occurs in breast cancer by investigating presence of DCX^+^/NFL^+^ cells in luminal-like and basal-like breast tumors. We generated an *in vitro* model by combining hESC-derived neural progenitor cells and breast cancer spheroids to mimic breast cancer innervation.

The communication between cancer cells and nerves is bidirectional^29^, by which cancer cells secrete neurotrophic factors to stimulate neural growth and infiltration, whereas neurotransmitters are released from nerve endings towards both cancer cells and surrounding stromal cells. Recent findings have demonstrated that direct neural and cancer cell contacts, not only via their neurotransmitters, are critical for tumor progression in breast cancer^8^. This supports the notion that investigating direct interactions between neural cells and breast cancer cells are necessary to understand the role and impact of tumor innervation. Experimental *in vitro* models suitable for investigating direct cancer-nerve interactions are limited. Previous studies have used murine neuron-like cell lines, dorsal root ganglia (DRG) and PC12 cells, to model cancer-nerve crosstalk^30, 31^, including a recent direct-contact co-culture study of DRG and TNBC cells highlighting a new role of sensory neurons in aggressive breast tumors^32^. Here we present a novel experimental co-culture model using *in vitro* differentiation of hESCs to generate human NPCs that maintain their multipotency in co-culture with breast cancer spheroids. We were unable to reproduce the findings from the PyMT-MMTV breast cancer model, claiming that multipotent NPCs can give rise to mature sympathetic neurons in the TME^5^. DCX+ cells in patient breast cancer tissue, did not express tyrosine hydroxylase (TH), nor DCX+ cells in our *in vitro* co-culture model by proteomics analysis. Although, we did detect TH expression during *in vitro* differentiation of NPCs into dopaminergic neurons at week 2 and week 5 following midbrain differentiation. Further studies are needed to determine whether NPCs in the TME of breast cancer holds the potential to generate mature and functional sympathetic neurons.

The bidirectional interplay between neural progenitors and breast cancer cells is rarely studied. Our *in vitro* model of NPCs and breast cancer spheroids enables monitoring of how breast cancer cells and NPCs interact in real time while allowing subsequent characterization of cell proteomes before and after co-culture. A previous study of MDA-MB-231 and the brain metastatic breast cancer cell line (MDA-MB-231Br) co-cultured with NPCs demonstrated boosted proliferation of MDA-MB-231Br in co-culture, while MDA-MB-231 fail to proliferate, with NPCs differentiating into astrocytes^33^. Our proteomics data indicated that co-culture with NPC induced aggressive tumor features in MDA-MB-231, such as induction of proteins involved in cell proliferation, migration, and invasion. At the same time, the proteome of NPCs showed downregulation of astrocyte markers such as GFAP and glutamine synthase when co-cultured with MDA-MB-231.

We found that the stromal compartments of breast tumors both contain DCX+ cells. Some of the positively labeled fibers appeared to be nucleated, an observation also found in prostate tumors^5^ and observed previously in innervated head and neck squamous cell carcinomas^7^, using other neural markers. Whether these cells are CNS-derived neurons, rather than sprouting axons in breast tumor stroma, remains to be elucidated.

In prostate cancer, the reported DCX expression was strongly associated with histological grade, and clinical outcome, by using immunofluorescent staining to identify stromal DCX^+^ cells^5^. A more recent report, evaluating DCX expression measured by transcriptome-wide microarray data of whole tissue from radical proctectomy specimen^34^, did not find significant differences between normal prostate, primary prostate cancer, and metastases, and found no increase with histological grade in a larger patient cohort. Whole tissue analysis could be limited by the heterogenous contributions from both stroma and tumor cells in the analyzed samples. We here show that stroma of breast tumors, of both luminal-like and basal-like phenotype, contain DCX+ cells that are also expressing other markers for neural progenitor cells. Our IMC analysis of breast tumors we separated stromal from epithelial tissue compartments prior to quantification of DCX+ cells, the proportion of stromal DCX+/NFL+ cells were higher in larger tumors and with higher histological grade and basal-like breast cancer phenotype. As the cohort size is limited, this warrants further assessments of neural progenitors in breast cancer tumorigenesis.

Our hESC-derived NPCs were able to integrate with both basal-like (MDA-MB-231, HS578T, BT549) and luminal-like (MCF7, BT474, T47D) breast cancer spheroids, but at varying degrees. Although these differences, likely caused by the morphological differences observed in basal-like and luminal like cells during spheroid formation, could indicate a preference of NPC integration in basal-like spheroids, our data does not hold sufficient statistical power to determine such breast cancer subtype specific differences. Previous studies have linked high nerve density to lymph node metastasis and poor patient outcome, but without clear differences in breast cancer subtypes^28, 35^.

Combined, we identified DCX+/NFL+ cells in breast tumors, and the presence of stromal DCX+/NFL+ cells was higher in larger tumors with higher histological grade in our cohort of 107 breast cancer cases, and the proportion of stromal DCX+/NFL+ appeared to be higher in basal-like breast cancer from our single-cell analyses. We established a new direct-interaction co-culture model mimicking breast cancer NPC-innervation, and proteomics analysis of these co-cultures corroborated that NPC interactions induces aggressive tumor features in breast cancer. Considering the stromal presence of DCX+/NFL+ cells, we believe that our model may serve as a useful tool for future studies to investigate how neural progenitors and sprouting axons interacts directly with other stromal cells of the breast cancer TME, such as immune and endothelial cells.

## Methods

### Breast cancer patient cohorts

Breast cancers from two independent study cohorts were included in our analysis. Cohort 1 comprises from a collection of primary invasive breast carcinomas diagnosed in women aged 50-69 as part of the prospective Norwegian Breast Cancer Screening Program in Hordaland County, Norway, during 1996-2003^36–38^. For this cohort, 41 tumor tissues were assessed using the available tissue microarrays (TMAs). The TMAs used were constructed as previously described in^37^.From them nine were luminal A, 14 luminal B and 18 triple negative, basal-like breast cancers. Cohort 2 is a BRCA case-control series established in cooperation with Prof. William D. Foulkes, McGill University, Canada^38, 39^ collected from patients counselled at the Hereditary Cancer Clinic at the McGill University Health Centre (MUHC) and the Jewish General Hospital, Montreal, Canada during 1981-2005. Cohort 2 was supplemented from a selection of confirmed BRCA negative cases from our archive at Haukeland, University Hospital and tested by the Department of Genetics, Haukeland University Hospital as part of a previous project. For this cohort, 76 tumor tissues were assessed using the available TMAs, with 37 being BRCA1 germline mutation positive and 39 BRCA mutation negative cases. Clinicopathological information for both cohorts (n=117) can be found at (Supplementary Table 2). Our focus in this report was on basal-like and luminal-like breast cancer (n=107), and we therefore did not include IMC analyses of the 10 HER2+ cases included in the cohort.

### Imaging mass cytometry (IMC)

<u>Antibody panel:</u> The IMC panel comprised 35 lanthanide (Ln) metal-conjugated antibodies (Supplementary Table 1), in addition to the free metals: iridium which binds double stranded DNA, and ruthenium which binds to tissue components^40^. Half of the antibodies were purchased pre-conjugated (Fluidigm) including CD45, CD3, CD4, CD8, FoxP3, CKAE1/AE3, and CD31. The remaining antibody clones, herein CD34 and Stathmin, were conjugated in our lab as described below. CD31 and CD34 were conjugated to the same metal isotope.

<u>Conjugation of antibodies:</u> The antibodies were conjugated to their respective metal isotope using the Maxpar® X8 antibody labeling kits and protocol (Fluidigm, CA, USA). Five µL of working solution of the metals and 100 ug of the glycerol- and carrier-free antibodies were used for conjugation as described in the protocol. The quantity of the conjugated antibody was determined using a NanoDrop spectrophotometer measuring absorbance of the conjugate at 280 nm. All antibodies were eluted with 20 µL W-buffer (Fluidigm) and diluted to 0.5 mg/ml (or 1:1 for the weakest antibodies) with antibody stabilizer (Candor Biosciences, Wangen, Germany) and stored at 4°C.

<u>Antibody validation:</u> All antibodies included in our panel were validated by IHC. Stains were performed on a test-TMA with positive control tissues such as tonsillar tissue, placenta, hippocampus, cerebellum, autonomic ganglion and peripheral nerve tissue, normal breast tissue, and selected breast carcinomas (ER+/PR+/HER2+; ER-/PR-/HER2- and basal-like tumors). In some cases, IHC staining patterns from the Human Protein Atlas ^41^ were used for comparison. As part of the validation process, IMC test stains were performed “in-house” on the test-TMA and a pilot-TMA (n=10) with four luminal-like, one HER2 positive, and five basal-like breast carcinomas.

<u>IMC staining protocol:</u> Antibody hybridization was performed according to the “Imaging Mass Cytometry Staining Protocol for FFPE Sectioned Tissue” (Fluidigm) with slight modifications. Freshly cut TMA slides underwent dewaxing, rehydration, and antigen retrieval for 48 minutesin a Ventana Discovery Ultra Autostainer (Roche Diagnostics GmbH) using CC1-buffer (pH 9). Slides were washed with a soap detergent and rinsed in hot water to remove the oil before they were transferred to a Coplin jar and washed with Maxpar H^2^O and then MaxPar phosphate buffered saline (PBS) (Fluidigm). To avoid non-specific binding, slides were blocked with 3% freshly made bovine serum albumin (BSA)/PBS buffer (Sigma-Aldrich/Merck). The antibody mix containing the individually diluted metal-conjugated antibodies in 0.5% BSA/PBS was applied to the slides, and the slides were stored overnight at 4°C in a hydration chamber. After antibody incubation, slides were washed first in 0.2% Triton X-100/PBS (Thermo Scientific) and then in Maxpar PBS before being stained with 0.3 µM Iridium (Ir)-intercalator (Fluidigm) for 30 minutes. Next, slides were washed in Maxpar H^2^O and then incubated with 0.0005% Ruthenium (RuO^4^)/PBS (Electron Microscopy Sciences)^40^ for 3 minutes. Finally, the slides were briefly washed in Maxpar H_2_O and air-dried.

<u>IMC analysis and data pre-processing:</u> Data from 117 breast cancer cases were acquired by a Helios time-of-flight mass cytometer (CyTOF) coupled to a Hyperion Imaging System (Fluidigm). The square inscribed in each circular TMA core (diameter 1.0 mm) was laser ablated at 200 Hz at a resolution of approximately 1 µm^2^. Pre-processing of raw data was performed using the CyTOF Software (v7.0.8493; Fluidigm), while the MCD™ Viewer software v1.0.560.6 (Fluidigm) was used for visualization of IMC images. The ImcSegmentationPipeline was utilized to process the raw data for downstream analyses^42, 43^. Using histoCAT (v1.7.6)^44^, the marker intensities and the spatial and morphological features of all images were exported in the csv-file format. To note that despite a well-performed segmentation, nuclei-mismatched signals might occur, especially in highly cellular areas. Such nuclei-mismatched signals appear when signals from “overlapping cell units” that do not capture the nucleus of an individual cell, are assigned to neighboring cells. Inspired by the methodology presented by Keren and colleagues^45^, cell type annotation was performed in a hierarchical scheme using unsupervised clustering and prior knowledge of cell type defining markers of our antibody panel. The deployed procedure operates in two steps: initially, single cells are categorized as immune or non-immune. FlowSOM algorithm^46^ was used to cluster the data into 120 clusters, that were further merged based on cluster cosine similarity. After this step, the annotated images were visually inspected by expert pathologists, and compared with the corresponding HE images. Areas with low quality within the tissues, like necrosis, scarring, or other tumor features, were flagged resulting to seven cases that were processed separately from the rest of the cohort. After this inspection step, Phenograph algorithm^47^ was used to refine the immune vs. non-immune annotation and split immune cells into three groups (B cells, T cells, macrophages), and non-immune cells into three groups (epithelial cells, endothelial cells, stromal cells). Lastly, the same procedure was repeated one more time to divide T cells to three subgroups namely CD8 TILs, CD4 TILs (CD4 positive, FoxP3 negative), and FoxP3 TILs (CD4 positive, FoxP3 positive), as well as epithelial cells to basal (CK5/6 and/or CK14 positive), luminal (non-basal, CK8/18 positive), and others (remaining epithelial cells expressing CKAE1/AE3 but no other keratins). (Supplementary Figure 1).

<u>Downstream analysis of neural markers using IMC:</u> After the single-cell annotation step, we used R language to investigate the intensity level of the neural markers, DCX, NFL, NCAM, INA and Nestin. We deployed a semi-automated gating strategy based on FlowDensity software^48^, a supervised clustering algorithm based on density estimation to define four groups of expression namely: DCX-NFL-, DCX-NFL+, DCX+NFL-, DCX+NFL+ (Supplementary Figure 1). Similarly, density estimations were used to define cells to High and Low groups based on the antibody intensity values of NCAM, INA and Nestin.

### Neural progenitor cell differentiation and passaging

Human embryonic stem cells, H9inGFPhES cells, (passage 37-40, WiCell Research Institute) were cultured in mTeSR1 medium supplemented with 0.5% Penicillin Streptomycin on Matrigel® coated wells. “Monolayer Culture Protocol” from “Generation and Culture of Neural Progenitor Cells Using the STEMdiff Neural System” (STEMCELL Technologies) was used to generate third passage NPC-like cells. Differentiation of Neural Progenitor Cells derived from XCL-1 DCXp-GFP were conducted in accordance with “Protocols for Neural Progenitor Cell Expansion and Dopaminergic Neuron Differentiation” from ATCC.

### Tumor cell spheroid generation and maintenance

The six basal- and luminal-like cell lines were disassociated with 0.25% trypsin and added individually to wells of a 6 well Ultra-low attachment plate in 2 mL Mammocult culturing media (STEMCELL Technologies). Accutase (STEMCELL Technologies) was further used to disassociate NPC cells. NPC were added to breast cancer cell containing wells equal to 20% of the total breast cancer cell number. Plates were kept on an orbital shaker (INFORS HT Celltron) at 70 rpm in a 37°C incubator for the duration of the co-culture. Media was changed every other day. Doxycycline (2 ug/mL) was added to the media to induce GFP expression in the NPCs 48 and 24 hours before collection. After 7 days of co-culture, spheroids were collected for processing. All cells used tested negative for mycoplasma contamination using MycoAlert Mycoplasm Detection Kit (Lonza, LT07-318).

### Fluorescence staining and imaging

<u>Whole mounting and coverslip:</u> Live spheroids were collected and fixated for 15 minutes in 10% formalin. To permeabilize the cells, spheroids were put into PBS with 2% Triton 100X for 2 hours. Protein blocking was done with 3 % BSA for two hours. The primary antibodies guinea pig polyclonal anti-doublecortin antibody (Merck, AB2253, 1:200 dilution), mouse monoclonal anti-CKAE1/AE3 (Dako, M3515, 1:200 dilution) were added to their respective spheroids and incubated for 2 days at 4 degrees. The following secondary antibodies were added at 1:500 concentration to their respective primary antibodies: Goat anti-guinea-pig AF 594 (A11073), donkey anti-rabbit AF 594 (A21207) & donkey anti-mouse AF 647 (A31571). Counterstaining was done with DAPI at a 1:1,000 solution for 24 hours at 4 degrees. Image acquisition and analysis was conducted with the Andor Dragonfly confocal microscope and Imaris 9.1.3 (Bitplane AG). All confocal tile-scan images were merged as a maximum projection.

<u>Neural cells:</u> Imaging was conducted using the same protocol principles as for the spheroids. However, neural cells were collected on coverslips before processing. Following the whole mounting protocol, neural cells were stained with rabbit polyclonal anti-tyrosine hydroxylase antibody (Merck, AB152, 1:200 dilution), guinea pig polyclonal anti-DCX antibody (Merck, AB2253, 1:200 dilution), rabbit polyclonal anti-INA antibody (Merck, AB5354, 1:200 dilution), mouse monoclonal anti-NeuN antibody (Merck, MAB377, 1:200 dilution), mouse monoclonal anti-Nestin antibody (CST, 33475, 1:200 dilution), rabbit monoclonal anti-Ki-67 antibody (Epredia, RM9106S, 1:200 dilution), chicken polyclonal anti-NFH antibody (Merck, AB5539, 1:200 dilution). The following secondaries were utilized for neural cell and breast cancer tissue staining: goat anti-guinea pig secondary antibody AF488 (A11073), donkey anti-mouse, secondary antibody AF594 (A21203), donkey anti-rabbit secondary antibody AF594 (A21207), goat anti-rabbit secondary antibody AF647 (A27040), goat anti-chicken secondary antibody AF647 (A21449), donkey anti-mouse secondary antibody AF647 (A31571). Slides were then mounted and imaged using a Leica TCS SP8 STED 3X confocal microscope.

<u>Breast cancer tissue</u>: For immunofluorescent staining, breast cancer tissues were serially sectioned at 4 μm. Xylene and graded ethanol washes were then used to deparaffinize and rehydrate the tissue. Tris/EDTA pH9 antigen retrieval was used to expose antigen binding sites. Non-specific binding was blocked in a 3% BSA containing PBS buffer. Slides were stained with mouse monoclonal anti-CKAE1/AE3 antibody (Dako, M3515, 1:200 dilution), guinea pig polyclonal anti-doublecortin antibody (Merck, AB2253, 1:200 dilution), chicken polyclonal anti-NFH antibody (Merck, AB5539, 1:200 dilution) overnight at 4 degrees. For secondary staining, antibodies AF488 (A11073), AF594 (A21203) AF647 (A21449) and a 1:1000 DAPI solution for 1.5 hours at RT. Slides were then mounted and imaged using a Leica TCS SP8 STED 3X confocal microscope.

### Histology and hematoxylin-eosin staining

Spheroids were fixated in 10% formalin for 15 minutes, washed with PBS twice and resuspended into a 5% agarose in PBS solution. The agarose solution containing spheroids was quickly centrifuged to center spheroids before the agarose block was cast in paraffin. Sections of 4 μm was cut from the paraffin block. Sections were subsequently deparaffinized and rehydrated using xylene and graded ethanol washes, respectively. Hematoxylin and eosin (H&E) staining was then performed using standard procedures.

### Spheroid FACS sorting

Spheroids were dissociated using Accutase (STEMCELL Technologies), incubated on shaker table for 10 minutes at 37°C before being manually pipetted until a single cell suspension remained. Cells were washed with PBS before being transferred to a PBS solution with 5% FBS. 5 minutes before sorting, cells were stained 1:500 with propidium iodide.

## Proteomics

### Cell lysis, protein digestion and Single-pot, solid-phase-enhanced samples preparation

Cells were washed in PBS and spun down before the cell pellets were resuspended in lysis buffer containing an 8 M Urea, 200 mM EPPS pH8.5 and protease inhibitors (Roche Complete with EDTA (Sigma Aldrich), homogenized, and sonicated in water bath for 30 seconds, three times. Protein concentration was determined by a Qubit 3.0 Fluorometer. In total we performed four TMT16-plex experiments (n=4). After lysis, hESC-derived neural cells were reduced with 100 mM DTT (DiThioThreitol, CAT# 171318-02, Amersham Biosciences) for 1h at RT, alkylated with 200 mM IAA (Iodoacetamide, CAT# I-6125, Sigma Aldrich) for 1h at RT (in the dark), followed by digestion of proteins into peptides using Trypsin Porcine (Promega, CAT# V5111) at trypsin-to-protein ratio 1:50 at 37 °C on a shaker. Reaction was quenched with formic acid. Breast cancer cells were processed using Single-pot, solid-phase-enhanced sample preparation (SP3) protocol with Sera-Mag™ SpeedBead Carboxylate-Modified [E3] Magnetic Particles (Thermo Scientific, CAT#65152105050250, CAT#45152105050250) at a 10:1 (wt/wt) bead/protein ratio^49^. Samples were desalted with Oasis Elution plates. Plates were prepared by centrifuging 500 μL of 70% acetonitrile (ACN) & 1% formic acid (FA) at 200g for 1 min. After, 500 μL 5% ACN & 1% FA were centrifuged twice at 200g for 1 minute. Samples were added and centrifuged at 100g for 3 min. 500 μL 5% ACN & 1% FA were added and centrifuged thrice at 200g for 1 min each. For elution, 100 μL 70% ACN & 1% FA were added twice at 100g for 3 min. Peptides were subsequently concentrated in a SpeedVac.

### Tandem Mass Tag (TMT) 16-plex labelling

TMTpro reagents (0.8 mg) were dissolved in anhydrous acetonitrile (40 μL) of which 7 μL was added to the peptides (50 µg) with 13 μL of acetonitrile to achieve a final concentration of approximately 30% (v/v). Following incubation at room temperature for 1 h, the reaction was quenched with hydroxylamine to a final concentration of 0.3% (v/v). TMT-labeled samples were pooled at a 1:1 ratio across all samples. For each experiment, the pooled sample was vacuum centrifuged to near dryness and subjected to C18 solid-phase extraction (SPE) (Sep-Pak, Waters).

### Off-line basic pH reversed-phase (BPRP) fractionation

We fractionated the pooled, labeled peptide sample using BPRP HPLC^50^ and an Agilent 1200 pump equipped with a degasser and a UV detector (set at 220 and 280 nm wavelength). Peptides were subjected to a 50-min linear gradient from 5% to 35% acetonitrile in 10 mM ammonium bicarbonate pH 8 at a flow rate of 0.6 mL/min over an Agilent 300Extend C18 column (3.5 μm particles, 4.6 mm ID and 220 mm in length). The peptide mixture was fractionated into a total of 96 fractions, which were consolidated into 12 super-fractions^51^, from which we analyze non-adjacent superfractions (n=12). Samples were subsequently acidified with 1% formic acid and vacuum centrifuged to near dryness. Each super-fraction was desalted via StageTip, dried again via vacuum centrifugation, and reconstituted in 5% acetonitrile, 5% formic acid for LC-MS/MS processing.

### Liquid chromatography and tandem mass spectrometry

Mass spectrometry data were collected using an Orbitrap Eclipse mass spectrometer (Thermo- Fisher Scientific, San Jose, CA) coupled to a Proxeon EASY-nLC 1200 liquid chromatography (LC) pump (ThermoFisher Scientific, San Jose, CA). Peptides were separated on a 100 μm inner diameter microcapillary column packed with ∼40 cm of Accucore150 resin (2.6 μm, 150 Å, ThermoFisher Scientific, San Jose, CA). For each analysis, we loaded ∼2 μg onto the column and separation was achieved using a 90min gradient of 6 to 28% acetonitrile in 0.125% formic acid at a flow rate of ∼420 nL/min. For the high-resolution MS2 (hrMS2) method, the scan sequence began with an MS1 spectrum (Orbitrap analysis; resolution, 60,000; mass range, 400−1600 Th; automatic gain control (AGC) target 100%; maximum injection time, auto). All data were acquired with FAIMS using three CVs (-40V, -60V, and -80V) each with a 1 sec. TopSpeed method. MS2 analysis consisted of high energy collision-induced dissociation (HCD) with the following settings: resolution, 50,000; AGC target, 200%; isolation width, 0.7 Th; normalized collision energy (NCE), 36; maximum injection time, 120 ms.

### Proteomics Data analysis

Mass spectra were processed using a Comet-based software pipeline^52, 53^. Spectra were converted to mzXML using a modified version of ReAdW.exe. Database searching included all entries from the human UniProt database. This database was concatenated with one composed of all protein sequences in the reversed order. Searches were performed using a 50-ppm precursor ion tolerance for total protein level profiling. TMTpro tags on lysine residues and peptide N termini (+304.207 Da) and carbamidomethylation of cysteine residues (+304.207 Da) were set as static modifications, while oxidation of methionine residues (+15.995 Da) was set as a variable modification. Peptide-spectrum matches (PSMs) were adjusted to a 1% false discovery rate (FDR)^54, 55^. PSM filtering was performed using a linear discriminant analysis, as described previously^56^, while considering the following parameters: XCorr, ΔCn, missed cleavages, peptide length, charge state, and precursor mass accuracy. For TMT-based reporter ion quantitation, we extracted the summed signal-to-noise (S/N) ratio for each TMT channel and found the closest matching centroid to the expected mass of the TMT reporter ion. PSMs were identified, quantified, and collapsed to a 1% peptide false discovery rate (FDR) and then collapsed further to a final protein-level FDR of 1%. Moreover, protein assembly was guided by principles of parsimony to produce the smallest set of proteins necessary to account for all observed peptides. Proteins were quantified by summing reporter ion counts across all matching PSMs, as described previously^56^. PSMs with poor quality and reporter summed signal-to-noise ratio less than 100, or no MS3 spectra were excluded from quantification^57^.

Correlation plot was made in Agilent GeneSpring GX software (version 14.9). Statistically significant proteins were identified by using t-tests between all cancer cell lines compared to co-cultured cancer cell lines (BH adjusted p < 0.05) with baseline transformation being applied to the median of all samples. Subsequent analysis was conducted using unbiased hierarchical cluster analysis (Spearman centered distance metrics with Ward’s drawing rules).

MDS, volcano plots and heatmaps were produced in R language using RStudio. For heatmaps, statistically significant proteins were identified by using t-tests between all cancer cell lines compared to co-cultured cancer cell lines (BH FDR, at a corrected p < 0.05). Volcano plots were generated by identifying significant proteins between MCF-7 or MDA-MB-231 and their co-cultured cancer cell line counterparts (BH FDR, at a corrected p < 0.01).

Significant proteins were imported into Pathway Ingenuity Pathway Analysis program (IPA®, QIAGEN Redwood City, www.qiagen.com/ingenuity), working as previously described^58^. In brief, following settings were used: Expression Fold Change (Exp Fold Change), Relationships to consider (Direct and Indirect Relationships), Reference set (Corresponding data analysis), Interaction networks (70 molecules/network; 25 networks/analysis), Molecule & Canonical Pathway subcategories were determined by “all” data types if not otherwise stated.

The underlying codes and programs implemented for this study may be available upon reasonable request to the corresponding author.

The mass spectrometry proteomics data have been deposited to the ProteomeXchange Consortium via the PRIDE repository with the dataset identifier: PXD040662.

### Ethics declaration

All experiments of this study have been approved by the Regional Committee for Medical and Health Research Ethics (2014/1984/REK Vest), as well as the work on human embryonic stem cells for generation of different neural cell types (212824/REK Vest).

## Acknowledgements

We thank Bendik Nordanger for excellent technical assistance. The confocal imaging was performed at the Molecular Imaging Center (MIC), Department of Biomedicine, University of Bergen. This work was partly supported by the Research Council of Norway through its Centre of Excellence funding scheme, project number 101651184. The study was also supported by the Norwegian Cancer Society, Regional Health Trust Western Norway (Helse Vest) and the Meltzer Research Fund and National Institutes of Health (NIH)/NIGMS grant R01 GM132129.

## Author contributions

OVB contributed with study design, cell culture work and data collection, analysis and interpretation of data and in writing the manuscript. MCF contributed with cell culture work and data collection and interpretation of data. KF and IW contributed with IMC data. DK contributed with analysis and interpretation of IMC data. CA, JBA and GK contributed with generating the tissue microarrays, collection of tissue and patient data. JAP contributed with MS-based proteomics analysis. LAA contributed with study design, histological evaluation of breast tumor tissue and interpretation of data. HV contributed with study design, writing the manuscript and interpretation of data. All authors have read, revised, and approved the manuscript.

