## Supplementary figures and images for "Global and single-cell proteomics view of the co-evolution between neural progenitors and breast cancer cells in a co-culture model"

### supp_figure1

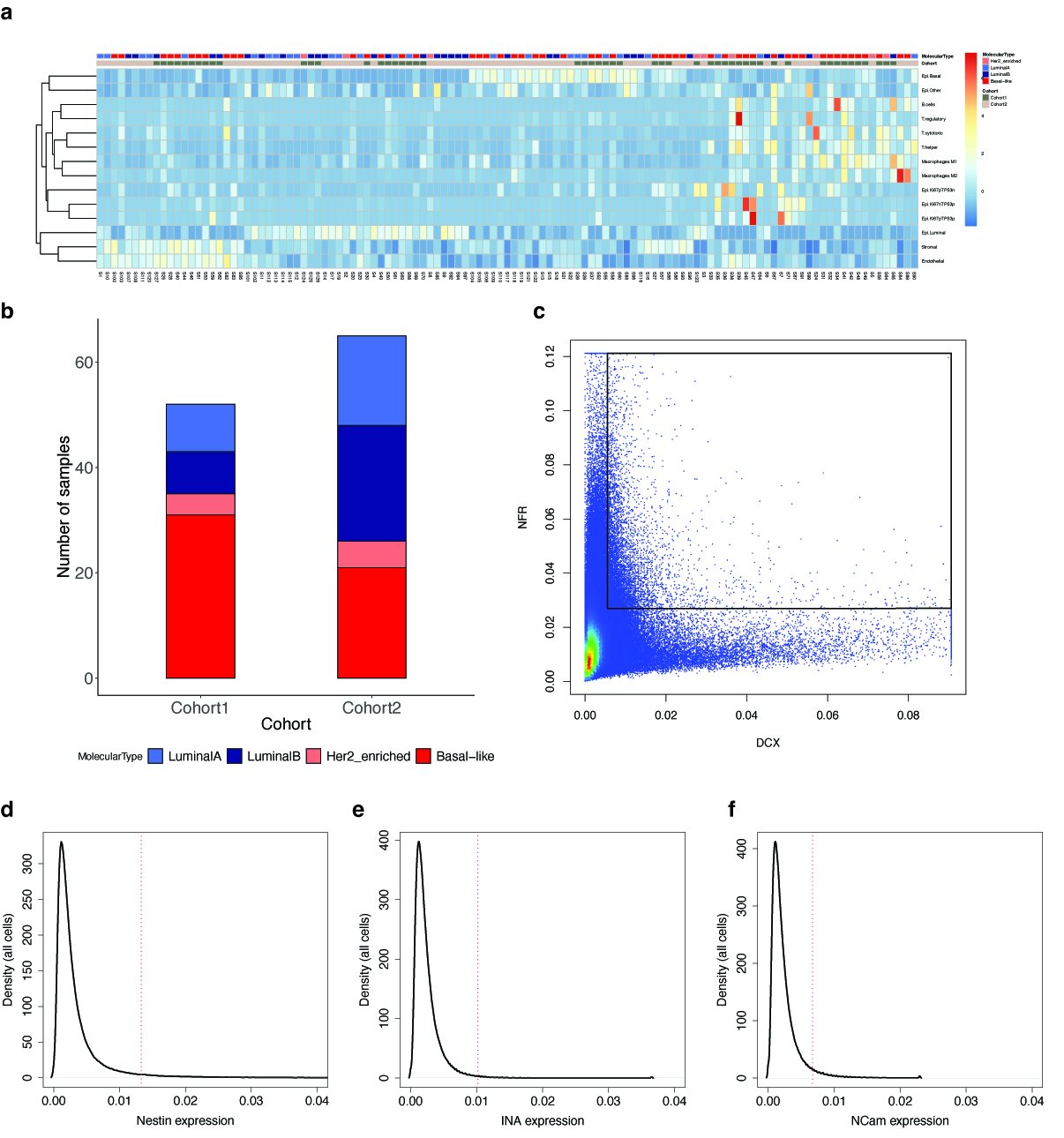
