## Supplementary material for "Global and single-cell proteomics view of the co-evolution between neural progenitors and breast cancer cells in a co-culture model": supp_table1

| **Cell type** | **Antibody** | **Description** |
| --- | --- | --- |
| Epithelial | CKAE1/AE3  CK5/6  CK14  CK8/18 | Cytokeratin (Pan CK)  Basal CK  Basal CK  Luminal CK |
| Stromal | alpha-SMA  Vimentin | Myofibroblast/Mesenchymal |
| Endothelial | CK31  CK34 | Endothelial |
| Immune | CD45  CD4  CD8  CD3  FoxP3  CD20  CD163  CD68 | Leukocyte (Pan immune)  T-cell  T-cell  T-cell  Regulatory T-cell  B-cell  Macrophage  Macrophage |
| Neural | Neurofilament-L (NFL)  Internexin-alpha (INA)  Doublecortin (DCX)  NCAM  Nestin | Neural progenitor  Neural  Neural and endothelial |
| Receptors/ Transcription factors | ER  PR  HER2  TP53  GATA3 | Estrogen receptor  Progesteron receptor  Human Epidermal Growth Factor Receptor 2  Tumor suppressor  Transcription factor |
| Miscellaneous | Histone H3  PD1, PDL-1, Foxa1, Ki67, Stathmin, PDGFR-beta, CD44, PDPN |  |

Supplementary Table 1
