## Supplementary material for "Global and single-cell proteomics view of the co-evolution between neural progenitors and breast cancer cells in a co-culture model": supp_table2

| # | Molecular subtype | Tumor Diameter | Histological Grade | Lymph node metastases |
| --- | --- | --- | --- | --- |
| 1; 4; 6; 7; 9; 10; 11; 12; 13; 14; 15; 16; 17; 18; 19; 20; 21; 22; 23; 25; 26; 27; 28; 29; 30; 31; 32; 33; 34; 35; 36; 37; 39; 40; 41; 42; 43; 44; 45; 46; 47; 48; 49; 50; 51; 52; 53; 54; 55; 56; 57; 58; 59; 60; 61; 62; 65; 66; 67; 68; 69; 70; 71; 82; 83; 84; 85; 86; 87; 88; 89; 90; 91; 92; 93; 94; 95; 96; 97; 98; 99; 100; 101; 102; 103; 104; 105; 106; 107; 108; 109; 110; 111; 112; 113; 114; 115; 116; 117; 118; 119; 120; 121; 122; 125; 126; 127 | Luminal A; Luminal B; Luminal B; Luminal B; Luminal B; Luminal A; Luminal B; Luminal A; Basal-like; Luminal B; Luminal B; Luminal A; Luminal A; Luminal B; Luminal A; Basal-like; Basal-like; Luminal A; Luminal A; Basal-like; Luminal A; Basal-like; Basal-like; Luminal A; Basal-like; Basal-like; Basal-like; Basal-like; Basal-like; Luminal A; Basal-like; Basal-like; Basal-like; Basal-like; Basal-like; Basal-like; Basal-like; Luminal A; Basal-like; Basal-like; Basal-like; Basal-like; Basal-like; Luminal B; Basal-like; Basal-like; Basal-like; Luminal A; Luminal A; Basal-like; Basal-like; Basal-like; Luminal B; Luminal A; Luminal A; Luminal B; Luminal B; Basal-like; Basal-like; Luminal A; Basal-like; Luminal B; Basal-like; Basal-like; Basal-like; Basal-like; Luminal B; Basal-like; Basal-like; Luminal B; Basal-like; Luminal A; Basal-like; Luminal B; Basal-like; Luminal B; Basal-like; Luminal B; Luminal B; Luminal B; Luminal B; Basal-like; Luminal B; Luminal A; Basal-like; Basal-like; Basal-like; Basal-like; Luminal B; Luminal B; Luminal A; Basal-like; Luminal A; Luminal A; Luminal A; Luminal B; Luminal A; Luminal B; Luminal B; Basal-like; Basal-like; Luminal A; Luminal A; Basal-like; Luminal B; Luminal B; Luminal B | 1; 0; 0; 0; 0; 0; 0; 0; 1; 1; 1; 1; 0; 0; 0; 1; 0; 0; 0; 1; 0; 1; 0; 0; 0; 0; 0; 1; 0; 0; 0; 1; 1; 0; 0; 0; 1; 0; 1; 1; 0; 1; 1; 0; 0; 0; 0; 1; 1; 0; 1; 0; 0; 1; 0; 1; 1; 0; 1; 0; 0; 0; 0; 1; 0; 1; 1; 1; 1; 0; 0; 0; 0; 1; 1; 0; 0; 1; 1; 1; 0; 0; 1; 0; 0; 0; 0; 0; 0; 0; 0; 0; 0; 1; 0; 1; 1; 1; 1; 0; 0; 0; 0; 1; 0 | 1; 3; 3; 2; 2; 1; 2; 2; 3; 2; 3; 1; 2; 3; 1; 2; 3; 1; 1; 3; 2; 2; 3; 2; 3; 3; 3; 3; 2; 3; 2; 3; 3; 1; 3; 3; 3; 3; 3; 3; 3; 2; 2; 1; 1; 3; 3; 2; 3; 2; 2; 3; 3; 3; 3; 1; 2; 2; 3; 3; 3; 2; 2; 2; 3; 3; 3; 2; 3; 1; 1; 3; 2; 2; 1; 2; 3; 3; 2; 1; 3; 2; 3; 2; 1; 1; 1; 2; 1; 1; 1; 2; 1; 2; 2; 3; 3; 2; 1; 3; 2; 3; 2 | 0; 0; 0; 0; 0; 0; 1; 1; 1; 1; 1; 1; 0; 1; 0; 0; 1; 0; 0; 1; 0; 0; 0; 0; 0; 0; 0; 1; 1; 0; 0; 1; 0; 0; 0; 0; 0; 0; 1; 0; 0; 0; 0; 1; 0; 0; 0; 0; 0; 1; 0; 0; 0; 0; 0; 1; 0; 0; 1; 0; 1; 0; 1; 0; 0; 0; 1; 0; 0; 0; 0; 1; 1; 0; 0; 0; 0; 1; 0; 0; 1; 1; 0 |
