## Supplementary material for "Global and single-cell proteomics view of the co-evolution between neural progenitors and breast cancer cells in a co-culture model": supp_table5

| | Breast cancer cells | | | | | | | | After co-culture | | | | | | | |
|---|---|---|---|---|---|---|---|---|---|---|---|---|---|---|---|---|
| **Gene symbol** | **Sample 1** | **Sample 2** | **Sample 3** | **Sample 4** | **Sample 5** | **Sample 6** | **Sample 7** | **Sample 8** | **Sample 1** | **Sample 2** | **Sample 3** | **Sample 4** | **Sample 5** | **Sample 6** | **Sample 7** | **Sample 8** |
| AHNAK2 | 11.3092831 | 11.2531443 | 9.99514529 | 9.8157035 | 11.2485028 | 11.0096467 | 10.0167108 | 9.9145122 | 10.9373223 | 10.9933489 | 10.6673156 | 10.5795988 | 12.2093346 | 12.1692392 | 12.0210042 | 10.2591665 |
| AKT1 | 8.28426426 | 8.36254441 | 8.20286825 | 9.12273742 | 8.58830238 | 8.37828185 | 8.98325301 | 9.2636401 | 8.20183477 | 8.10311992 | 8.30275772 | 8.17265252 | 8.74269343 | 8.76310279 | 7.85013057 | 8.02554379 |
| CASP8 | 7.98135607 | 8.16851167 | 9.02387353 | 8.47920226 | 7.24434481 | 7.33709557 | 9.03903692 | 8.28961779 | 8.22028158 | 8.42253577 | 8.24411167 | 8.34310575 | 8.78763162 | 8.78561267 | 7.63238456 | 7.86547347 |
| CDH1 | 9.58585452 | 9.33748142 | 9.56902202 | 9.35500965 | 10.5436892 | 10.5069918 | 9.94771583 | 10.003799 | 7.31855185 | 7.2134445 | 7.8756695 | 7.53556311 | 7.41102789 | 7.41740529 | 8.58548463 | 7.71245134 |
| CDH2 | 3.51120235 | 4.10994554 | 3.55952852 | 3.95861071 | 3.97714252 | 3.72363501 | 5.3652971 | 4.15605678 | 3.32231757 | 3.03082513 | 6.37534358 | 6.55352946 | 3.59106899 | 3.54260475 | 6.27271749 | 5.54332884 |
| DPYSL2 | 11.2082222 | 11.1306993 | 11.2500141 | 11.2846217 | 11.8788901 | 11.8186341 | 13.2627717 | 12.9935081 | 11.4684866 | 11.4031079 | 11.4072784 | 11.3598374 | 12.9053193 | 12.9636818 | 13.4079183 | 13.7892178 |
| DPYSL5 | 9.18721596 | 9.1411492 | 8.63772978 | 8.23779792 | 9.39857483 | 10.9646441 | 8.75784333 | 8.66399921 | 10.8203784 | 10.62411 | 10.1532607 | 10.8101217 | 8.94079809 | 8.90022379 | 11.5197198 | 8.50549004 |
| GIT1 | 7.9382505 | 7.80305263 | 8.03858433 | 7.69165978 | 7.56585608 | 7.71045519 | 7.79434438 | 7.54126702 | 7.0664552 | 6.64822095 | 7.99324412 | 7.37598643 | 7.78314231 | 7.69540231 | 6.71115071 | 6.87860274 |
| GRB2 | 11.2479453 | 11.2566864 | 10.496694 | 11.762764 | 10.7240875 | 10.7850192 | 10.0703088 | 10.2773688 | 9.80642395 | 9.96511761 | 9.82185209 | 10.8242348 | 10.3503094 | 10.3571561 | 9.67893459 | 9.80408261 |
| MAP2K4 | 8.54892207 | 8.65759714 | 9.00345045 | 8.75612261 | 7.67503257 | 8.17180729 | 8.63802314 | 8.5336451 | 8.70332569 | 8.70541813 | 8.62719825 | 8.8502306 | 8.91880064 | 8.86390129 | 8.06726225 | 8.26365887 |
| MUC1 | 9.38609313 | 9.48920774 | 9.24520803 | 9.15278171 | 10.3063352 | 10.7057274 | 9.49072054 | 10.0745964 | 9.22612073 | 9.44433577 | 9.43254608 | 9.30921257 | 11.0126315 | 11.0014363 | 11.784459 | 8.89678951 |
| MUC5B | 5.37001428 | 5.57803901 | 6.96897776 | 6.70463684 | 6.41693599 | 5.82499183 | 6.23962717 | 6.64107343 | 5.93765041 | 5.77877103 | 5.77946986 | 5.86479034 | 8.3829309 | 8.33428223 | 6.97108366 | 6.03769629 |
| PTPN11 | 11.6331178 | 11.5838397 | 11.82118 | 11.6034734 | 10.9814603 | 11.4174412 | 11.4701708 | 11.4281822 | 11.4999356 | 11.4846165 | 11.1264851 | 10.9594804 | 11.7421957 | 11.7815318 | 10.7817203 | 11.1596506 |
| PXN | 7.84984918 | 7.78909017 | 7.74323893 | 8.20172697 | 8.67990229 | 8.480875 | 7.75735667 | 7.92799124 | 8.80138826 | 8.56729445 | 8.22436833 | 8.03363774 | 8.56166218 | 8.67713117 | 8.34849359 | 8.17054102 |
| RAC1 | 10.4120193 | 10.2946552 | 12.0216775 | 11.4618733 | 9.57939703 | 9.71649368 | 10.9825872 | 10.0914089 | 10.9257102 | 11.0327896 | 10.5616984 | 10.6821343 | 11.0990848 | 10.9355923 | 9.39028721 | 9.96592855 |
| RTN4 | 8.94197486 | 8.80851698 | 8.35389225 | 8.32822163 | 9.08120646 | 9.35269233 | 8.60118765 | 8.9563355 | 8.93757097 | 9.21276873 | 9.41093676 | 8.70097716 | 10.0396731 | 9.99326393 | 9.8032335 | 9.61004114 |
| SHANK2 | 7.26189746 | 7.32755264 | 7.10989332 | 7.98297077 | 8.05586807 | 8.23089468 | 7.77178656 | 8.6712049 | 6.67649309 | 6.71304965 | 7.00811484 | 7.10436811 | 6.30892327 | 6.28213025 | 6.53361785 | 6.14098924 |
| TGFBR2 | 5.44573001 | 5.20064559 | 5.6759655 | 5.34782154 | 6.73665899 | 6.76875335 | 5.50651626 | 6.1932257 | 5.56505789 | 5.35781941 | 5.43349698 | 5.54299159 | 7.21302643 | 7.25987505 | 6.35174492 | 5.37661337 |
