## Supplementary material for "Global and single-cell proteomics view of the co-evolution between neural progenitors and breast cancer cells in a co-culture model": supp_table6

| | MCF-7 | | | MCF-7 after co-culture | | |
|---|---|---|---|---|---|---|
| **Gene symbol** | **Sample 1** | **Sample 2** | **Sample 3** | **Sample 1** | **Sample 2** | **Sample 3** |
| ABI1 | 6.37504638 | 6.31942465 | 6.48887969 | 6.05193323 | 5.95108884 | 5.92691268 |
| AHNAK | 13.4875512 | 13.5710771 | 13.6489979 | 12.9804377 | 13.0444505 | 12.9938123 |
| AHNAK2 | 7.87954037 | 7.74613806 | 7.92866641 | 8.15235076 | 7.9966825 | 8.17639758 |
| AKT1 | 8.54797029 | 8.66092291 | 8.54563251 | 8.84973348 | 8.77309304 | 8.96992752 |
| ARHGAP35 | 7.85650023 | 7.70116782 | 7.63911637 | 7.75374492 | 7.78011321 | 7.72723978 |
| BRAF | 2.82645697 | 3.25575601 | 3.51203721 | 3.63407046 | 3.36265223 | 3.69219283 |
| BRCA1 | 2.94288041 | 2.73102245 | 2.86860578 | 2.5439264 | 2.76406797 | 2.64710264 |
| BRCA2 | 3.53068256 | 3.13309822 | 3.52511672 | 3.37210238 | 3.5402629 | 2.92373489 |
| CALD1 | 6.8906534 | 6.7954293 | 6.66851629 | 6.94656712 | 7.07907939 | 6.96462967 |
| CASP8 | 7.469536 | 7.53782409 | 7.42628992 | 7.33013707 | 7.32242394 | 7.34341671 |
| CDC42 | 4.62851699 | 4.0120412 | 5.00204087 | 3.42977663 | 4.25541009 | 3.90558196 |
| CDH1 | 7.19698073 | 7.19846469 | 7.05526074 | 7.39309009 | 7.21519322 | 7.32944655 |
| CDH11 | 4.92948804 | 4.65612449 | 5.44565388 | 5.12045148 | 5.02301133 | 5.16905511 |
| CLPP | 6.82883465 | 6.75914251 | 6.75134321 | 7.28404212 | 7.20880986 | 7.09873746 |
| CTNNA2 | 8.7483304 | 8.74227238 | 8.39549129 | 8.29024007 | 8.35454472 | 8.37799566 |
| CTNNB1 | 7.29195343 | 7.56670101 | 7.33820154 | 6.85658738 | 6.9310093 | 7.10950683 |
| EGFR | 5.97042805 | 5.84335831 | 5.60556545 | 5.3612455 | 5.68444287 | 5.75312848 |
| ESR1 | 3.5854313 | 3.75025275 | 3.68574957 | 5.44297003 | 5.28768283 | 5.43885172 |
| FASN | 11.4541142 | 11.4540628 | 11.3852477 | 12.0084811 | 11.9915466 | 12.056465 |
| FGFR1 | 2.78067335 | 2.73086381 | 3.35931032 | 3.40232653 | 3.05468701 | 3.17263952 |
| FGFR1OP | 4.31101623 | 4.83815005 | 4.77706722 | 4.41860826 | 4.68399956 | 4.49274409 |
| FGFR1OP2 | 4.8883049 | 4.84168636 | 4.82485948 | 4.87005889 | 4.87350317 | 4.58230911 |
| KRAS | 5.28439938 | 5.13427845 | 5.30868487 | 5.03208321 | 4.86534974 | 5.03207439 |
| MAP2K4 | 6.68382275 | 6.60680493 | 6.59584973 | 6.00363147 | 6.26791028 | 6.38353323 |
| MAPT | 7.51341179 | 7.57731552 | 7.72977027 | 7.08164854 | 7.04624002 | 6.85325965 |
| MKI67 | 9.6849502 | 9.44995748 | 9.50485275 | 8.91981975 | 8.8238194 | 8.78845081 |
| MTOR | 7.62568693 | 7.52796955 | 7.46897424 | 7.18040649 | 7.26030799 | 7.33672077 |
| MUC1 | 6.18943913 | 6.1367943 | 6.20664322 | 7.15935672 | 7.16638405 | 7.33647978 |
| MUC5B | 6.41028343 | 6.44846656 | 6.44186854 | 6.49943135 | 6.49546653 | 6.52909942 |
| NRAS | 7.05905238 | 6.7600213 | 7.02321099 | 7.04138537 | 7.07303828 | 6.94184578 |
| NUP93 | 8.39315875 | 8.39345481 | 8.30503115 | 8.55326449 | 8.58528181 | 8.6737684 |
| PAK4 | 7.82435418 | 7.91163766 | 7.97936231 | 7.7735019 | 7.85632592 | 7.84017366 |
| PKP2 | 6.47745391 | 6.38726775 | 6.38257907 | 6.42262323 | 6.23944387 | 6.34019372 |
| PTEN | 6.55044513 | 6.57022611 | 6.5285135 | 6.32908771 | 6.32809684 | 6.19536925 |
| PTPN11 | 8.1152882 | 8.14978776 | 8.12036749 | 7.98031669 | 7.96314462 | 7.89249425 |
| PXN | 7.03907258 | 7.17373709 | 6.98235477 | 6.5870739 | 6.70419416 | 6.31726788 |
| RAC1 | 6.27373034 | 6.37700225 | 6.43526354 | 5.25103355 | 5.31957097 | 5.35914161 |
| RB1 | 7.94978587 | 7.7554017 | 8.02641278 | 7.98395695 | 7.88642211 | 7.8869159 |
| RHOA | 3.5598711 | 3.76582114 | 4.13419629 | 4.43741419 | 3.82614168 | 3.44201457 |
| RIPK2 | 3.99276843 | 4.1996959 | 4.17038979 | 4.52950239 | 3.9414444 | 4.07654199 |
| ROBO1 | 2.80944943 | 3.0630421 | 2.98556965 | 3.87797977 | 3.52560537 | 2.9079405 |
| ROCK1 | 7.4768191 | 7.36293446 | 7.70477515 | 7.63761025 | 7.67647194 | 7.55279204 |
| STAG2 | 7.23502813 | 7.30372596 | 6.98044808 | 7.4424874 | 7.50677964 | 7.52587312 |
| TGFB1 | 3.54799265 | 3.03607776 | 2.94391008 | 3.93929241 | 3.36567854 | 3.81562666 |
| TGFB1I1 | 4.13431953 | 4.04193507 | 4.45186389 | 4.20368667 | 4.20829094 | 4.325962 |
| TGFB2 | 4.31898199 | 4.68382556 | 4.09599191 | 4.63590527 | 4.04315199 | 4.51918962 |
| TGFBI | 5.14484175 | 5.13602729 | 4.89727916 | 5.08291118 | 5.18739958 | 5.29851536 |
| TGFBR2 | 3.93480897 | 3.70492728 | 3.42690879 | 3.37072228 | 3.37282676 | 3.73270335 |
| TGFBRAP1 | 6.36587797 | 6.23898743 | 6.19115992 | 6.85628857 | 6.89893387 | 6.8658446 |
| TP53 | 4.70790021 | 4.97370311 | 5.23140169 | 5.23259936 | 4.97853429 | 5.48492437 |
| TRIM21 | 6.3518721 | 6.52477519 | 6.24056048 | 6.62902578 | 6.38028096 | 6.4920182 |
| VIM | 10.7001536 | 10.6165856 | 10.7790545 | 10.8889411 | 10.8256584 | 10.8759643 |
