## Supplementary material for "Global and single-cell proteomics view of the co-evolution between neural progenitors and breast cancer cells in a co-culture model": supp_table7

| | MDA-MB-231 | | | MDA-MB-231 after co-culture | | |
|---|---|---|---|---|---|---|
| Gene symbol | Sample 1 | Sample 2 | Sample 3 | Sample 1 | Sample 2 | Sample 3 |
| ABI1 | 6.02015324 | 5.66049798 | 5.82101625 | 5.90201809 | 5.87326446 | 5.97053619 |
| AHNAK | 13.7660423 | 13.703471 | 13.7194423 | 14.7428673 | 14.6573408 | 14.6967557 |
| AHNAK2 | 9.24068402 | 9.16192798 | 9.17294998 | 10.747337 | 10.7455483 | 10.7368754 |
| AKT1 | 7.81001887 | 7.77202419 | 7.85433234 | 8.31256081 | 8.35841385 | 8.36206217 |
| ARHGAP35 | 7.44397954 | 7.5046045 | 7.62635152 | 7.80039519 | 7.82434781 | 7.81539611 |
| BRAF | 3.60314561 | 3.61176256 | 3.79491487 | 3.17729275 | 3.44128427 | 3.37992588 |
| BRCA1 | 3.45070979 | 3.45416273 | 3.27907008 | 2.70040642 | 2.92281629 | 2.44244726 |
| BRCA2 | 3.61620476 | 3.4231882 | 3.18352939 | 3.53953107 | 3.51008844 | 3.05642572 |
| CALD1 | 8.36541259 | 8.54552827 | 8.41638798 | 8.39008295 | 8.42634863 | 8.31583791 |
| CASP8 | 7.56027933 | 7.45777485 | 7.52355412 | 7.95218794 | 8.00740857 | 8.013278 |
| CDC42 | 4.68330906 | 4.15024283 | 4.29807173 | 4.20569747 | 3.82384105 | 3.92518293 |
| CDH1 | 5.63532198 | 5.36339539 | 5.61469218 | 5.3715275 | 5.35813952 | 5.77230534 |
| CDH11 | 6.51612391 | 6.3338441 | 6.47199379 | 7.52068939 | 7.44172415 | 7.49545533 |
| CLPP | 8.03494749 | 8.07136582 | 8.03741517 | 7.64310579 | 7.69046599 | 7.64310579 |
| CTNNA2 | 7.50248973 | 7.19781626 | 7.35972321 | 7.3431946 | 7.35312035 | 7.29851169 |
| CTNNB1 | 6.30650286 | 6.30884866 | 6.42549623 | 6.58843131 | 6.7351295 | 6.81125266 |
| EGFR | 8.04079382 | 7.71498042 | 7.72565937 | 7.36666317 | 7.32781309 | 7.22418502 |
| ESR1 | 3.38310981 | 3.3028352 | 3.3299703 | 3.31665697 | 3.5259561 | 3.60109031 |
| FASN | 10.8509682 | 10.8322334 | 10.8935746 | 12.1607601 | 12.1206299 | 12.0668595 |
| FGFR1 | 2.83379281 | 3.19976813 | 3.00082751 | 3.15757734 | 3.52541745 | 3.43978269 |
| FGFR1OP | 4.91853889 | 5.17720424 | 4.84431032 | 5.00991593 | 4.80529762 | 4.79653027 |
| FGFR1OP2 | 5.00743772 | 4.73341942 | 4.83205453 | 4.68177532 | 4.8958697 | 4.86035203 |
| KRAS | 5.04555655 | 5.00356626 | 5.12069615 | 5.113763 | 5.35392311 | 4.89995865 |
| MAP2K4 | 6.42143203 | 6.27321188 | 6.33929063 | 7.01171925 | 6.97859607 | 7.08022061 |
| MAPT | 3.97088126 | 3.75436376 | 3.83025532 | 4.75039212 | 4.36206304 | 4.44986507 |
| MKI67 | 8.37598209 | 8.34847146 | 8.33467106 | 8.28678414 | 8.19983427 | 8.29670977 |
| MTOR | 7.42488851 | 7.44466707 | 7.5936899 | 7.84629272 | 7.75232712 | 7.89819626 |
| MUC1 | 6.08143976 | 6.0181935 | 6.06265925 | 7.8115802 | 7.92879075 | 8.00273627 |
| MUC5B | 5.78744814 | 5.83094233 | 5.76932479 | 7.71013141 | 7.78468064 | 7.86899536 |
| NRAS | 5.94003281 | 5.4939643 | 5.72497762 | 5.84597923 | 5.65985037 | 5.52472505 |
| NUP93 | 8.71567377 | 8.70475787 | 8.67459129 | 8.49595493 | 8.38501249 | 8.39773024 |
| PAK4 | 7.49120374 | 7.38829288 | 7.52262076 | 7.69652248 | 7.39532848 | 7.55699645 |
| PKP2 | 6.70574296 | 6.54883732 | 6.59134282 | 7.27713859 | 7.21063238 | 7.16384085 |
| PTEN | 6.1525262 | 6.03935996 | 6.11331711 | 6.35360383 | 6.61603857 | 6.32598178 |
| PTPN11 | 8.0054449 | 7.99609493 | 8.00871433 | 8.36464246 | 8.36423098 | 8.30440152 |
| PXN | 7.18562872 | 7.23865313 | 7.12487607 | 7.44235469 | 7.55141604 | 7.50381779 |
| RAC1 | 6.00978832 | 6.17610531 | 6.05025136 | 6.41750825 | 6.35809907 | 6.28378661 |
| RB1 | 8.45107096 | 8.46151219 | 8.51082745 | 7.81711797 | 7.83873751 | 7.80012983 |
| RHOA | 4.32011638 | 4.59082355 | 4.6416674 | 4.73998848 | 4.98621067 | 4.84049345 |
| RIPK2 | 3.90378064 | 3.56900493 | 4.12272241 | 3.73038998 | 4.0074467 | 3.9683212 |
| ROBO1 231 | 2.6182457 | 1.78585182 | 2.70259992 | 3.21474341 | 2.49223264 | 2.35366204 |
| ROCK1 | 7.42193016 | 7.49417557 | 7.48601441 | 7.34573345 | 7.32186498 | 7.38123145 |
| STAG2 | 7.13676559 | 6.88634285 | 7.08252165 | 6.65643917 | 6.82137858 | 6.69221794 |
| TGFB1 | 4.32452261 | 3.87451174 | 4.28483243 | 3.85290773 | 4.03988412 | 4.28224704 |
| TGFB1I1 | 6.21967773 | 6.18987989 | 6.18950239 | 6.45651541 | 6.51448841 | 6.62047644 |
| TGFB2 | 5.58868466 | 5.42151955 | 5.71978581 | 5.13192691 | 4.99156266 | 4.73976171 |
| TGFBI | 5.88548407 | 5.80811986 | 5.81558824 | 4.86885465 | 4.33232125 | 4.41844634 |
| TGFBR2 | 3.86040666 | 3.89507961 | 3.82574493 | 6.18527389 | 6.24305971 | 5.96262768 |
| TGFBRAP1 | 6.38074157 | 6.26875813 | 6.16430559 | 6.309771 | 6.32782746 | 6.25128925 |
| TP53 | 8.59169064 | 8.43363968 | 8.42542993 | 7.53560199 | 7.58766502 | 7.54619995 |
| TRIM21 | 6.71046207 | 7.00424294 | 6.86272413 | 7.27241517 | 7.33891462 | 7.03182524 |
| VIM | 13.2853083 | 13.4471091 | 13.4111534 | 12.8033824 | 12.787368 | 12.7827505 |
